# *TLS2trees*: a scalable tree segmentation pipeline for TLS data

**DOI:** 10.1101/2022.12.07.518693

**Authors:** Phil Wilkes, Mathias Disney, John Armston, Harm Bartholomeus, Lisa Bentley, Benjamin Brede, Andrew Burt, Kim Calders, Cecilia Chavana-Bryant, Daniel Clewley, Laura Duncanson, Brieanne Forbes, Sean Krisanski, Yadvinder Malhi, David Moffat, Niall Origo, Alexander Shenkin, Wanxin Yang

## Abstract

Above Ground Biomass (AGB) is an important metric used to quantify the mass of carbon stored in terrestrial ecosystems. For forests, this is routinely estimated at the plot scale (typically ≥1 ha) using inventory measurements and allometry. In recent years, Terrestrial Laser Scanning (TLS) has appeared as a disruptive technology that can generate a more accurate assessment of tree and plot scale AGB; however, operationalising TLS methods has had to overcome a number of challenges. One such challenge is the segmentation of individual trees from plot level point clouds that are required to estimate woody volume, this is often done manually (e.g. with interactive point cloud editing software) and can be very time consuming. Here we present *TLS2trees*, an automated processing pipeline and set of Python command line tools that aims to redress this processing bottleneck. *TLS2trees* consists of existing and new methods and is specifically designed to be horizontally scalable. The processing pipeline is demonstrated across 10 plots of 7 forest types; from open savanna to dense tropical rainforest, where a total of 10,557 trees are segmented. *TLS2trees* segmented trees are compared to 1,281 manually segmented trees. Results indicate that *TLS2trees* performs well, particularly for larger trees (i.e. the cohort of largest trees that comprise 50% of total plot volume), where plot-wise tree volume bias is ±0.4 m^3^ and %RMSE is ^~^60%. To facilitate improvements to the presented methods as well as modification for other laser scanning modes (e.g. mobile and UAV laser scanning), *TLS2trees* is a free and open-source software (FOSS).

## 1. Introduction

Above Ground Biomass (AGB) is an important metric that quantifies the amount of carbon stored in terrestrial ecosystems, and as such has been identified as an Essential Climate Variable (ECV). However, accurate quantification of forest AGB is a significant and ongoing challenge owing to a number of factors, including systematic errors when applying allometry to inventory data (Burt et al., 2020). To improve the accuracy of AGB quantification in forests, groups such as the International Panel on Climate Change (IPCC) and the Committee on Earth Observing Satellites (CEOS) have identified Terrestrial Laser Scanning (TLS) as a disruptive technology (Ogle et al., 2019, Duncanson et al., 2021).

TLS is an active remote sensing technology that generates a detailed 3D point cloud of the surrounding area with centimetre to millimetre accuracy (Newnham et al. 2015, Calders et al. in Duncanson et al. 2021). A TLS-based approach has been used to estimate AGB across a range of forest types (Demol et al., 2022a), for example; tropical forests (Momo Takoudjou et al., 2018, Gonzalez de Tanago et al., 2018, Beyene et al., 2020, Burt et al., 2021, Levick et al., 2021, Brede et al., 2022), temperate forests (Calders et al., 2015, Stovall et al., 2017, Disney et al., 2020, Calders et al.), mangroves (Feliciano et al., 2014) and trees outside forests (Wilkes et al., 2018, Kükenbrink et al., 2021, Van Den Berge et al., 2021). Over the past decade, area scanned has increased from a few trees to systematic acquisition across multiple hectares that replicates forest inventory protocols (Wilkes et al., 2017).

The benefits of TLS based methods are that they are non-destructive, capture tree plasticity particularly of large trees (Burt et al., 2021) and are (mostly) free from errors and assumptions associated with allometric modelling (Chave et al. in Duncanson et al. 2021). When compared to destructive harvest, TLS has been shown to be more accurate than the application of existing allometries, particularly for larger trees that contribute disproportionately to plot-level AGB (Demol et al., 2022a). However, operationalising plot-level (i.e. ≥ 1 ha) TLS workflow as a “turn-key” solution to produce an AGB product has yet to be fully realised (Martin-Ducup et al., 2021).

A TLS survey of a forest plot to estimate AGB typically involves: (1) capturing scan data from multiple fixed positions across a plot, (2) co-register scans to produce a single plot-level point cloud *P*, (3) instance segmentation of *P* into a set of point clouds that represent individual trees *S*, (4) semantic segmentation of *s* ∈ *S* into wood and leaf point classes, (5) estimate of woody volume for *s* ∈ *S* e.g. using a Quantitative Structure Model (QSM) approach, and (6) conversion of volume to AGB via an estimate of wood basic density. Steps 1-2 have been largely been solved with improvements in scanner technology and standardised scanning protocols (Calders et al., 2020, Wilkes et al., 2017). There are also a number of existing methods for semantic segmentation (Vicari et al., 2019, Wang et al., 2020) and QSM generation, where *s* ∈ *S* are enclosed in a geometric primitives e.g. a set of cylinders (Raumonen et al., 2013, Hackenberg et al., 2015, Stovall et al., 2017). Conversion of volume to mass is an ongoing challenge for all non-destructive methods of AGB estimation as wood density varies greatly within and between trees and geographic regions (Phillips et al., 2019). Therefore, it is suggested, that from a TLS workflow perspective at least, the most significant remaining challenge to operational plot level AGB estimation is the instance segmentation in step 3 i.e. *S* ⊂ *P*.

*P* is encoded with geometric information of whole-tree structure, for all trees in the surveyed area regardless of size. If *P* is of sufficient density and quality then *S* can be accurately segmented and an unbiased assessment of plot-level AGB, with associated uncertainties, can be produced (Calders et al., 2015, Momo Takoudjou et al., 2018, Burt et al., 2021). Currently, the most accurate method of *S* ⊂ *P* is to manually segment individual trees from their neighbours and other vegetation using interactive point cloud editing software (Momo Takoudjou et al., 2018, Gonzalez de Tanago et al., 2018, Disney et al., 2020, Brede et al., 2022). Manually segmenting trees increases tree- and plot-level estimation accuracy by a factor of 10 and 3 respectively, compared to existing automated TLS pipelines (Martin-Ducup et al., 2021). However, manual segmentation can be very time consuming (10’s of minutes per tree) as well as subjective and difficult to reproduce or validate. These factors have limited the number (and provenance) of segmented trees, where total segmented trees is limited to 10’s (and only in rare cases 100’s) of trees per hectare.

Automated whole-tree instance segmentation methods have been previously demonstrated (Tao et al., 2015, Burt et al., 2019, Wang et al., 2021, Martin-Ducup et al., 2021). Tao et al. (2015) presented a method where a graph is constructed through clustered *P*, then using a Dijkstra shortest-path method (Dijkstra, 1959) clusters are mapped to a base node resulting in segmented *S*. They applied their method to TLS data collected in 3 forest patches and reported high accuracy for number of trees correctly identified and completeness of extracted trees. Subsequent methods have built upon these graph based methods and have been applied in different forest types such as Schelorphyll, coniferous and tropical forests (Wang et al., 2021, Martin-Ducup et al., 2021). Another method was presented by Burt et al. (2019) who used rule constrained clustering with proximity testing to segment trees in a tropical forest plot; this method also requires an allometric assumption of *DBH* to crown height and extent. These methods have shown promise in demonstration plots, this paper aims to build on these and present a method that is scalable and is applicable across forest types.

There are a number of challenges that can hinder achieving sufficient quality of *S* ⊂ *P* with an automated workflow (Demol et al., 2022a). These include factors attributable to scanning protocol and scanner specification, such as; scanner type and optics (Newnham et al., 2015, Calders et al., 2020), sufficient sampling density to minimise occlusion (Wilkes et al., 2017), co-registration accuracy and co-alignment errors (Demol et al., 2022b) and post-processing computational constraints e.g. large data volumes (data volumes are typically >65 Gb ha^-1^). Further, automated segmentation also needs to be sensitive to forest demography and composition where trees in a plot can range in size (height, diameter etc.) over orders of magnitude, neighbouring crowns may intersect, parasite species such as lianas could be present (Moorthy et al., 2019) and understorey vegetation maybe dense.

To address the interoperability and scalability challenges of automated tree segmentation from plot-level point clouds, we present *TLS2trees*. *TLS2trees* is a set of Python command line tools that are free and open-source software (FOSS) and are specifcially designed to be horizontally scalable (Kissling et al., 2022), for example, on a High Performance Computing (HPC) facility. The output is a set of segmented point clouds of individual trees, where points are classified into leaf and wood components. Below, *TLS2trees* is demonstrated at ten forest plots that cover seven forest types; from open savanna to dense tropical forest. Additionally, segmented point clouds are compared to a set of 1,053 manually segmented reference trees.

## 2. Materials and Methods

### 2.1. Data acquisition

The TLS data used here were captured from ten plots across seven different forest types (Table 1). At each plot, data were acquired with a RIEGL VZ-400 scanner (RIEGL Laser Measurement Systems GmbH, Horn, Austria) where a set of scans *I* were captured on a regular grid (Wilkes et al., 2017) or, for RUSH plots, in a star formation (Calders et al., 2015). At each scan position two scans were acquired where the scanner rotation axis was orientated perpendicular then parallel to the ground surface. Manually-placed reflectors were used as tie points between each scan position to aid co-registration. Post processing was done using RiSCAN Pro software (versions 2.1 - 2.9) where indvidual scans were co-registered to a common coordinate system on a per plot basis. Once registration was complete, a set of 4 × 4 transformation matrices *M* was exported for each scan.

**Table 1:**
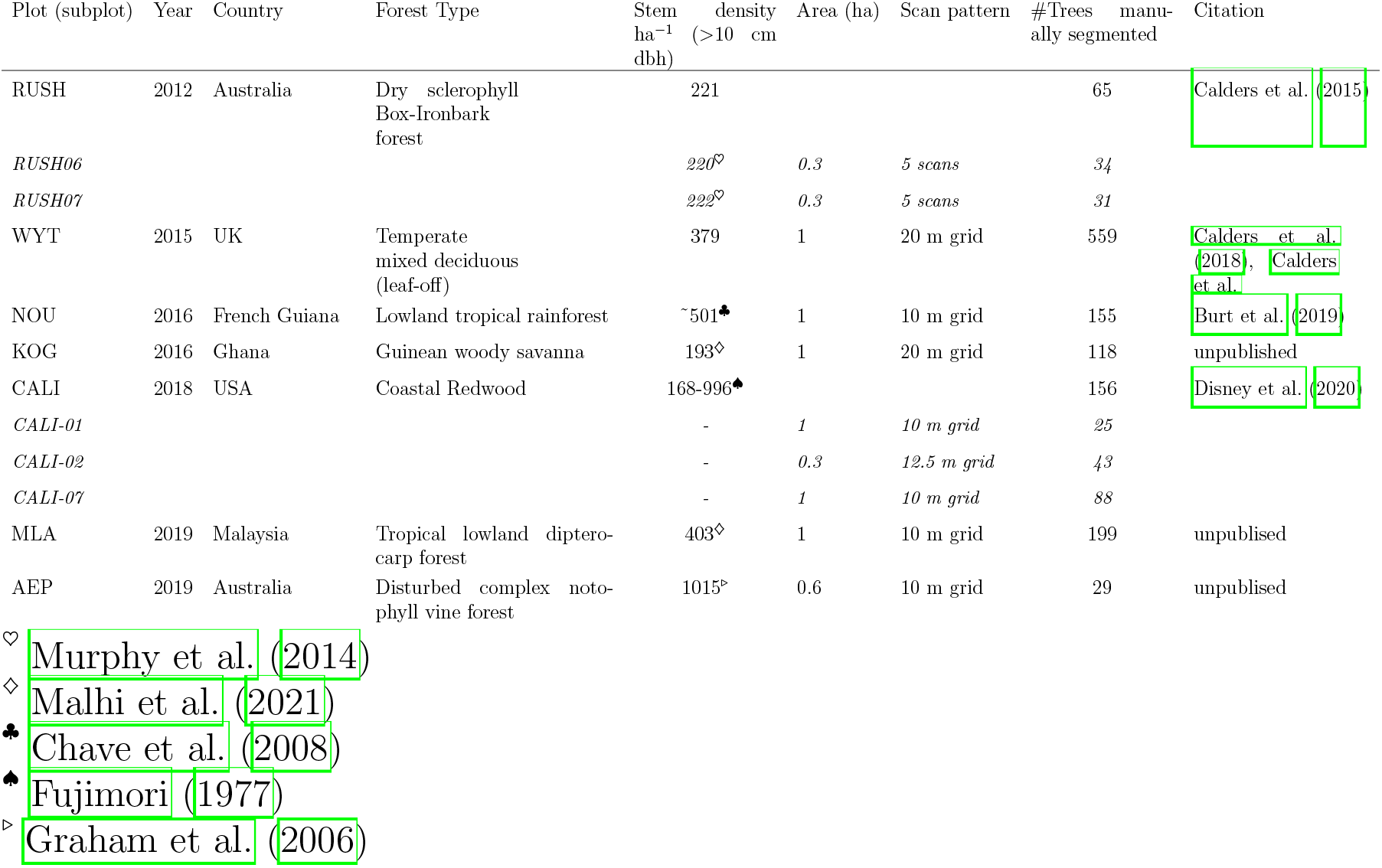
TLS plots.

### 2.2. TLS2trees

The *TLS2trees* software package consists of a set of Python command line tools. *TLS2trees* can be considered Free and Open Source Software (FOSS) and is licensed under Creative Commons BY 4.0 (https://creativecommons.org/licenses/by/4.0/). The workflow combines modified versions of previously published routines along with new methods that are described in detail below, it is also modular and therefore new or additional methods can be added or existing steps replaced. Processing is horizontally scalable i.e. can be scaled across multiple computing nodes (Kissling et al., 2022) where this is achieved using a tile-based workflow. For more information and code see https://github.com/philwilkes/TLS2trees.

This section presents a detailed description of the workflow. As is shown in Figure 1, a coregistered global point cloud *P* is passed through the *TLS2trees* pipeline to generate a set of segmented tree point clouds *S*. The pipeline is presented in three steps: (1) pre-processing (Section 2.2.1), (2) semantic segmentation where points are classified into different components (Section 2.2.2), and (3) an instance segmentation where *P* is segmented to a set of individual trees *S* (Section 2.2.3). Once *S* ⊂ *P*, structure traits, such as total woody volume, are computed via TreeQSM (Section 2.4). Exposed workflow parameters for each step are listed in the Appendix A.

**Figure 1:**
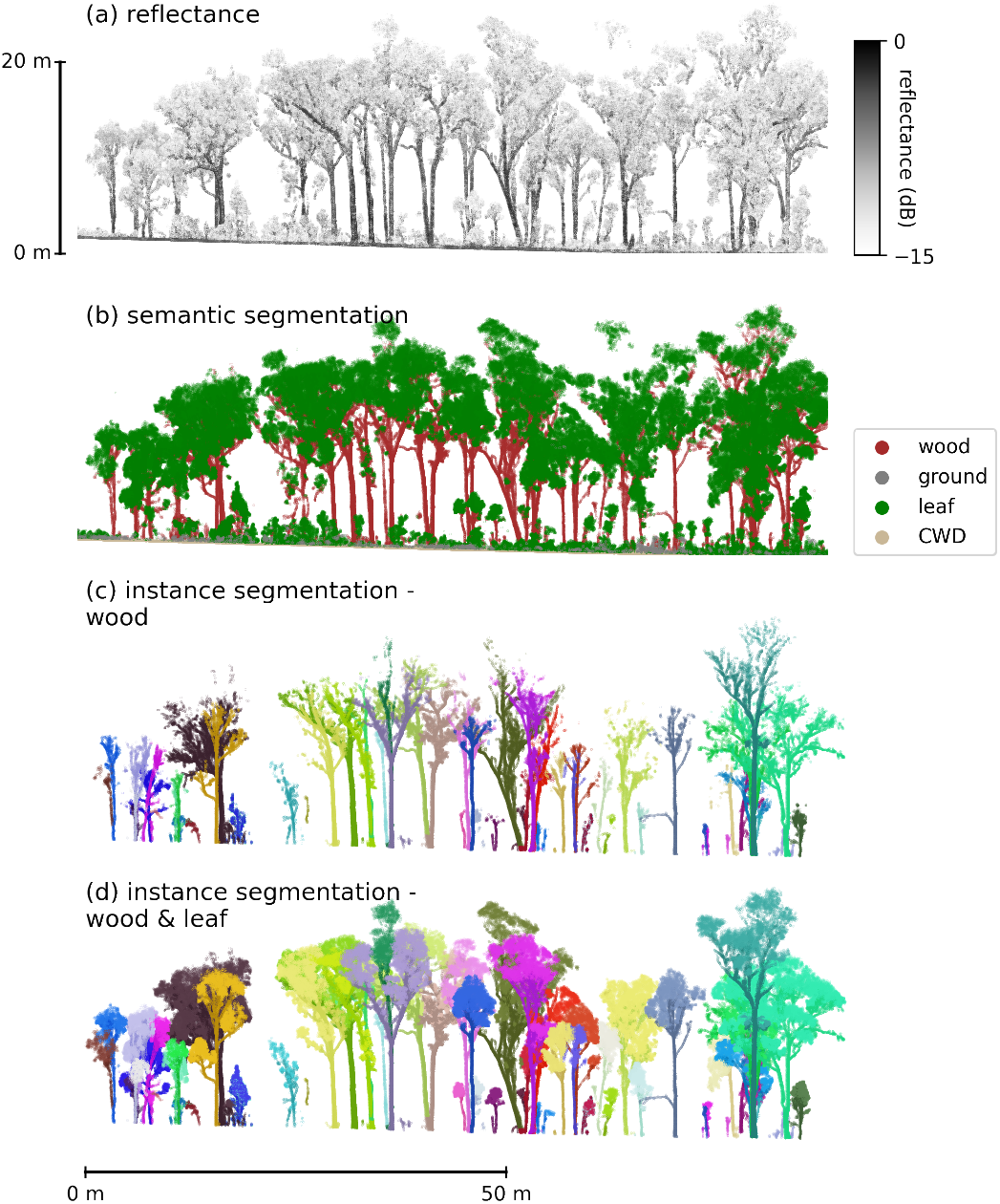
*TLS2trees* applied to a 70 m × 20 m strip of forest in plot RUSH where (a) *P* is coloured by calibrated reflectance (b) semantic segmentation into 4 classes using *FSCT* (Krisanski et al., 2021) (c) instance segmentation where only points classified as wood are displayed and segmented trees are coloured randomly, and (d) instance segmentation where leaf points are attributed to individual stems.

#### 2.2.1. Step 1. Preprocessing

The workflow starts at the point of a set of individual individual scans *I* and corresponding rotation matrices *M*; the first step is (*I, M*) ↦ *P*. During (*I, M*) ↦ *P, I* are clipped to the plot extent determined by M, plus a 10 m buffer (Martin-Ducup et al., 2021). P is then projected onto a 10 m × 10 m grid to produce a set of tiled point clouds *T* where individuals tiles *t* ∈ *T* are the processing unit for subsequent steps. A tile index of *T* is also generated to map the spatial location of neighbouring tiles.

As is inherent with TLS data, objects closer to the scanner are over sampled whereas objects further away (e.g. the top of the canopy, or at the edge of a scanned region) can be undersampled (Burt et al., 2019). To mitigate the impact of this difference and to reduce file size, *P* is downsampled to a common point density using PDAL’s *Voxel Center Nearest Neighbor* method (PDAL Contributors, 2020) where here a voxel edge length of 0.02 m is used.

#### 2.2.2. Step 2: Semantic segmentation

Semantic segmentation is the process of labelling points into homogeneous groups; here this is into classes of different biophysical components e.g. ground, leaf, wood, etc. Previous workflows have classified points simultaneously with or after instance segmentation (Wang, 2020, Vicari et al., 2019, Wang et al., 2020). However, instance segmentation can be hampered by the presence of leaves that reduce gaps between tree crowns, or causes neighbouring crowns to intersect.

Here, a modified version of the Krisanski et al. (2021) *FSCT* semantic segmentation method is applied prior to instance segmentation. *FSCT* uses the Pointnet++ deep-learning method (Qi et al., 2017) applied via Python’s PyTorch. Using a pre-trained model, P is classified into 4 classes; ground (G), woody (W), leaf (L) or coarse woody debris (X) (Figure 1b and equation 1).

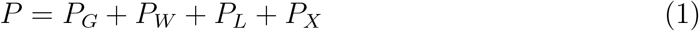

The semantic segmentation is applied to *t* + *t_b_* ∈ *T*, *b* refers to neighbouring tiles used to generate a 5 m buffer that mitigates any edge effect; once the semantic segmentation is complete only the class label for t are retained. It should be noted that the pre-trained model has not been modified since its initial release by Krisanski et al. (2021) i.e. the model is not trained for forest types specific to Table 1.

#### 2.2.3. Step 3: Instance segmentation

Instance segmentation is the process of identifying and segmenting individual trees *S* = {*s*_1_, *s*_2_,…, *s_k_*} encoded in *P* i.e. *P*_*W*+*L*_ ↦ *S*. Here, this is achieved in a two step process where (1) *P_W_* are grouped into a set of individual woody stems *S* (Figure 3C), then (2) *P_L_* are assigned to *s* ∈ *S* (Figure 3D).

**Figure 2:**
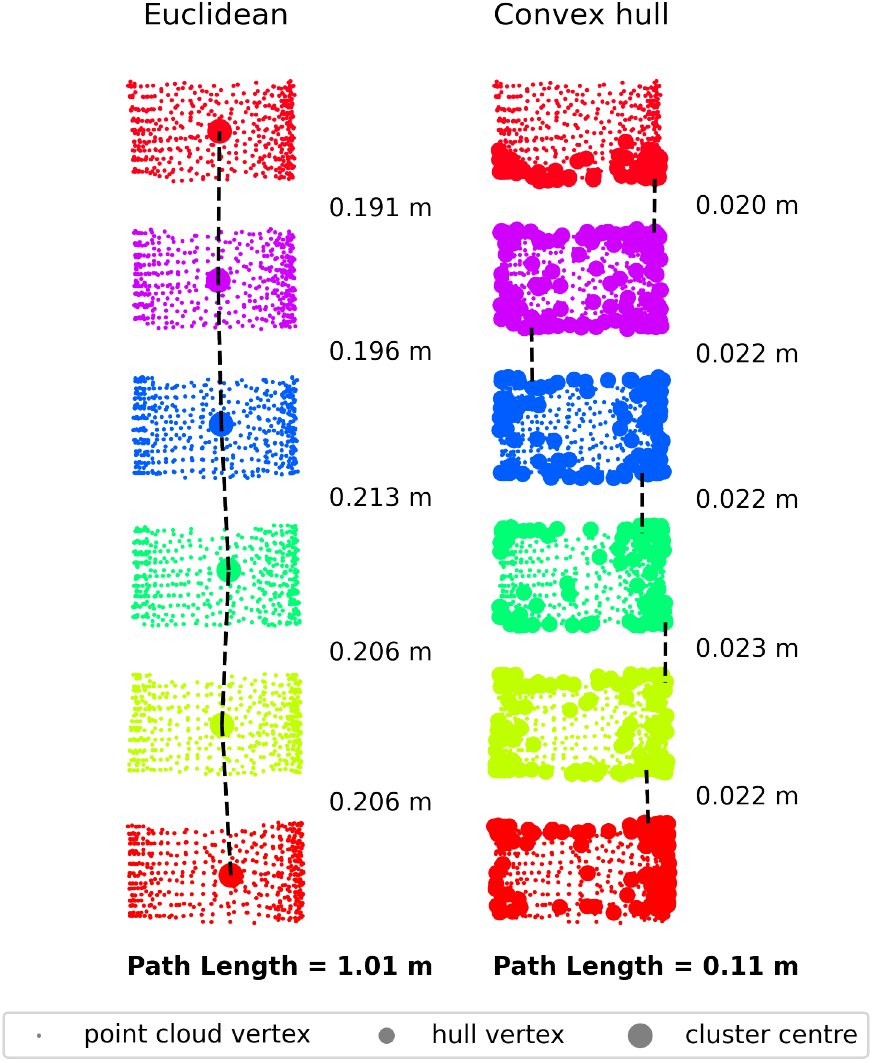
A comparison of methods for the construction of graphs through point clouds. Displayed is a point cloud section along a stem where adjacent clusters have been exploded so connections between clusters are clearly visible. For the “Euclidean” method, edge weights (values to the right) are calculated as the Euclidean distance between cluster centres; this is synonymous with path length (Tao et al., 2015). For the “Convex hull” method, edge weights are calculated as the distance between the vertices of neighbouring convex hulls (i.e. connectivity).

**Figure 3:**
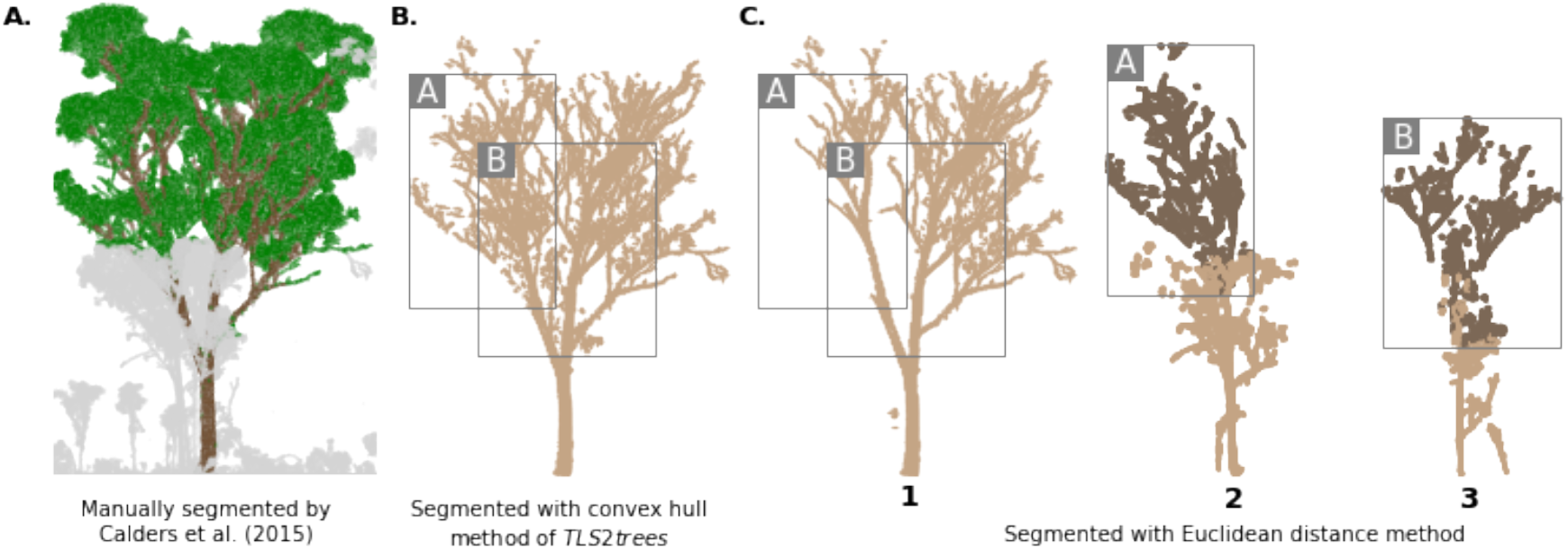
A tree segmented using (A) manual segmentation by Calders et al. 2015 (B) segmented using the convex hull method of *TLS2trees* and (C) segmented using a Euclidean distance method. Grey boxes A and B highlight branches that have been wrongly attributed to Trees 2 and 3 respectively (dark brown points in Panel C). Trees 2 and 3 (Panel C) can be seen at the base of the manually segmented tree (grey points in Panel A).

Both steps use a Dijkstra’s shortest path method (Dijkstra, 1959) where a graph *G* = (*N, E*) is constructed; *N* are a set of nodes and *E* are a set of edges that connect *N*. A path *p* can be defined that connects two nodes (*n, n_k_*) ∈ *N* where *p* = [*n, n*_1_, *n*_2_, … *n_k_*]; for each (*n,n_k_*) pair there are multiple solutions to *p*. To determine the shortest path between the pair (*n, n_k_*) a weight function 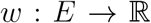 is defined (equation 2) (Tao et al., 2015).

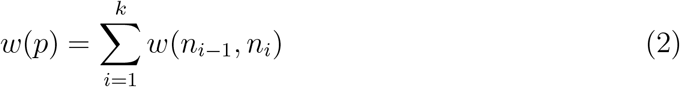

Previous methods have constructed graphs where *N* is a set of vertices in 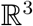 and *w* are the Euclidean distances between *N* (Tao et al., 2015, Wang et al., 2021, Vicari et al., 2019, Brede et al., 2022). Therefore the shortest path *min*{*w*(*p*) : *n* ↦ *n_k_*} is analogues to a vascular system (Tao et al., 2015) (Figure 2 left). However, experimentation here found that if one or more tree crowns intersect then this method results in a poor instance segmentation (Figure 3). To solve this, a different approach is taken to compute *w*.

To define a graph *G_W_* for wood classified points *P_W_*, a set of nodes *N_W_* are first generated. To do this, *P_W_* is sliced vertically (relative to ground normalised height calculated from *P_G_*) at intervals of 0.2 m; slicing is required to regularise the distance between clusters. Then for each slice, DBSCAN (Ester et al., 1996) is used to map *P_W_* to a set of clusters *C_W_* (Figure 2). DBSCAN parameters are dependent on acquisition and point cloud characteristics; after downsampling an *eps* value of 0.1 m and *minimum_sample* of 20 points are used here.

For each cluster *c* ∈ *C_W_*, a convex hull is computed generating a set of hull vertices *V_W_* where *V_W_* ⊂ *C_W_* ⊂ *P_W_* (Figure 2 right). A convex hull retains information on the occupation of space and proximity to neighbouring clusters, whereas collapsing a cluster to a single vertex does not. A k-nearest neighbour search is then performed on *V_W_* to identify vertices in neighbouring clusters; within cluster connections are disregarded. The Euclidean distance where *min*(*dist*(*V_W_c_x___*, *V_W_c_y___*)) is used as the edge weight function *w* (Figure 2 right). *w*(*p*) is therefore determined by distance between cluster edges (i.e. connectivity) as opposed to distance between cluster centroids (i.e. path length). A parameter is available to set allowable maximum distance between *V_W_c_x___* and *V_w_c_y___* (Appendix A).

Dijkstra shortest path analysis requires a subset of source nodes *n_b_* ⊂ *N_W_* from which to calculate distance from. Here *n_b_* are generated by taking a slice through *C_W_*[*z*_1_, *z*_2_] where *z*_1_ and *z*_2_ are upper and lower bounds of the slice relative to ground height. RANSAC cylinder fitting is then used to identify stem bases as opposed to noise such as leaves (Burt et al., 2019). It is important to attempt to identify *all* stems, regardless of diameter, otherwise smaller stems can be erroneously included into larger neighbouring stems; here a lower limit of ø = 0.05 m was used for stem detection.

Once the graph *G_W_* is generated, the Dijkstra shortest path method is used where multiple source nodes *n_b_* can be defined. The output of the shortest path analysis is a disjoint union of undirected acyclic graphs. Then *N_w_* ↦ *C_W_* and therefore *P_W_* can be mapped to a set of individual stems *S*.

Once step 1 is complete, another graph *G_L_* is created that maps leaf classified points *P_L_* (and *P_W_* that are not attributed to a stem) to *s* ∈ *S*. *N_L_* are computed by first mapping *P_L_* to a set of voxels *C_L_* with an edge length of 0.5 m; *C_L_* is equivalent to the clusters *C_W_* above. A set of 6 vertices are generated for *c* ∈ *C_L_* where a vertex is assigned to the centroid of each voxel facet; this results in *V_L_* which are equivalent to *V_W_* above. *E_L_* and corresponding w are determined by calculating the Euclidean distances to neighbouring *v* ∈ *V_L_*. A set of source nodes *n_b_* ⊂ *N_W_* are taken from *G_W_* where *n* ∈ *N_W_* have no children i.e. *n* is a branch tip; *n_b_* are already associated with *s* ∈ *S*. Shortest path analysis is again used to connect *N_L_* with *n_b_* and therefore assign *P_L_* to *s* ∈ *S*.

To run the instance segmentation, each *t* ∈ *T* is buffered by neighbouring tiles (see Table 2 for example buffer size). Once complete, *S* is pruned so that only trees whose germination point is within t are retained; further, a first-pass filter to remove trees where *DBH* <0.1 m (as determined by the RANSAC cylinder fitting) is applied.

**Table 2:** Processing steps and benchmark times.

|  | GPU/CPU | Parallelisable | Software | RUSH | NOU | MLA |
| --- | --- | --- | --- | --- | --- | --- |
| Plot area (ha) |  |  |  | 0.3 | 1 | 1 |
| Number of scans |  |  |  | 10 | 242 | 242 |
| Raw data size (Gb) |  |  |  | 2.4 | 65.7 | 107.8 |
| Preprocessing | CPU | Within machine<br>10x CPU used<br>for benchmark | Python, PDAL |  |  |  |
| Raw points per tile |  |  |  | 400K | 33.2M | 67.7M |
| Downsample points per tile |  |  |  | 350K | 5.5M | 7M |
| Wall time |  |  |  | 47 secs | 45 mins | 1 hr 20 mins |
| Semantic segmentation | GPU | TRUE | Python, PyTorch |  |  |  |
| Mean CPU time per tile |  |  |  | 1 min | 13 mins | 20 mins |
| Maximum RAM usage per tile (gb) |  |  |  | 4.6 | 39.7 | 82.3 |
| Wall time |  |  |  | 11 mins | 4 hrs | 3.5 hrs |
| Instance segmentation | CPU | TRUE | Python, NetworkX |  |  |  |
| Buffer (tiles) |  |  |  | 3 x 3 | 5 x 5 | 7 x 7 |
| Mean CPU time per tile |  |  |  | 16 mins | 3 hrs | 4 hr 50 mins |
| Maximum RAM usage per tile |  |  |  | 2.8 | 51.6 | 129.8 |
| Wall time |  |  |  | 1 hr 30 mins | 16 hrs | 18 hrs |

### 2.3. Manually segmented trees

A total of 1,281 trees have been manually segmented in previous studies (Table 1). Methods for selecting trees to manually segment from P differed for each plot. For RUSH plots, trees were selected by the Victorian State Government to update statewide AGB allometry, where a subset of trees were harvested across a range of sizes and species (Murphy et al., 2014, Calders et al., 2015). For WYT, all trees within a central 1 ha plot were segmented from a larger 6 ha scanned area, this reduces edge effects inherent at other plots. Individual tree point clouds were also split into >1 tree if bifurcation occurred <1.3 m (Calders et al., 2018). All trees were segmented from NOU where *DBH* >0.2 m (Burt et al., 2019) and KOG where *DBH* >0.1 m. For AEP and MLA plots, tree species that comprised 80% of total basal area were selected and a 2-3 trees from each species were segmented (Shenkin et al., 2020). It should be noted no trees have been specifically manually segmented or modified for this manuscript, i.e. trees were segmented before the inception of this method.

Trees from plots RUSH, MLA, AEP and KOG were manually segmented from the plot-level point cloud using either RiSCAN Pro or CloudCompare v2.X (https://www.danielgm.net/cc Data from WYT (Calders et al., 2018, Calders et al.), NOU (Burt et al., 2019) and CALI (Disney et al., 2020) were first segmented with *treeseg* (Burt et al., 2019), after which trees were modified manually e.g. removing overlapping crowns. During this process all trees were manually verified for commission and omission errors by an experienced operator.

After segmentation, all tree point clouds underwent a semantic segmentation into leaf and wood points using the *TLSeparation* Python package (Vicari et al., 2019); the exception being WYT where data were captured in leaf-off conditions. Using the wood classified point cloud only, per tree structural traits were then modelled using *TreeQSM* (see Section 2.4).

### 2.4. Quantitative Structure Models

Quantitative Structure Model (QSM) methods enclose the wood-only point clouds in a set of geometric primitives e.g. a cylinder. This allows for the estimation of morphological and topological traits such as volume, length and surface area metrics (Raumonen et al., 2013). *TreeQSM* (version 2.3.1, Raumonen 2019) is used to generate a QSM for all manually and automatically segmented trees. Since the version of *TreeQSM* and iterated parameter space may differ from previously published versions, it should be noted that modelled values for manually segmented trees may differ from previously published results.

A set of *TreeQSM* parameters control the overall fit of cylinders, following Raumonen et al. (2013) three parameters are iterated over here; *PatchDiam*1 = [0.20, 0.22, …, 0.3], *PatchDiam*2*Min* = [0.5, 0.7,…, 0.15] and *PatchDiam*2*Max* = [0.15, 0.17,…, 0.25]. *TreeQSM* is run in Octave (Eaton et al., 2020) where, for each parameter set permutation 5 models are generated. This results in a total of 625 models per segmented tree. An optimal model is then selected by minimising the point to cylinder surface distance (Burt et al., 2019, Martin-Ducup et al., 2021).

### 2.5. Comparing tree pairs

To assess the accuracy of segmentation, a corresponding pair of trees is identified in the manually and *TLS2trees* segmented sets. Pairs are identified by firstly taking a slice through all segmented trees between 2-3 m and computing the centroid of the slice. Tree pairs are then matched by selecting a tree from the *TLS2trees* data set that mostly closely matches the position of a manually segmented tree. Trees where more than one target tree is within 1 m of the reference tree or where a match is >2 m from the reference tree are disregarded from further analysis.

For matched tree pairs, QSM and point cloud metrics are compared. QSM metrics include total woody volume (m^3^), trunk volume (m^3^), total branch length (m) and *DBH* (m). Point cloud metrics include leaf-on crown height (m), leaf-on projected crown area (m^2^), wood/leaf point classification ratios and Jaccard Index metrics. The Jaccard Index (Jaccard, 1912) is a measure of spatial concordance; values range from 0-100% where 0% and 100% indicate no overlap and complete overlap of point clouds accordingly (Brede et al., 2022). A Weighted Jaccard Index is calculated here by first voxelising the segmented point clouds with an edge length of 0.5 m, a weight *w* is then assigned to each voxel corresponding to the number of points per voxel. Then for each tree pair the weighted intersection and union of voxel sets is computed using Python’s sklearn (Pedregosa et al., 2011) *jaccard_index* method.

Owing to the unbalance in number of segmented trees per plot, when deriving metrics for all matched pairs, a bootstrap sampling approach is taken where for each iteration a sample of 10 trees per plot is drawn.

### 2.6. Computing infrastructure and software

Insufficient computing infrastructure can be a bottle neck to processing large geospatial data sets, such as point cloud data. TLS data can generate particularly large data sets for relatively small regions of interest when compared to other laser scanning instruments e.g. airborne. Therefore, for a new TLS data processing pipeline to be operationalised it would ideally be capable of being horizontally scaled.

Horizontal scaling is achieved in the semantic and instance segmentation steps of *TLS2trees* by mapping *P* to a set of tiles *T*; the *t* ∈ *T* are run independently and in parallel. To process the plots in Table 1, the UK’s Natural Environment Research Council’s (NERC) JASMIN computing facility and the NERC Earth Observation Data Acquisition and Analysis Service (NEODAAS) MAssive GPU for Earth Observation (MAGEO) cluster were used. The JASMIN facility (https://jasmin.ac.uk/) is designed for data intensive computing and comprises a large volume of storage combined with general purpose batch computing capability. The MAGEO cluster (https://www.neodaas.ac.uk/) is a specialised system designed for earth observation data and comprises 40 GPUs, 200 GPU cores and 0.5 PB of fast storage. Data processing requirements and times for a subset of plots are presented in Table 2

All steps use Python as a base programming language, this includes common scientific libraries such as Numpy, Scipy and Pandas (see requirements.txt for full list) which are managed with the *conda* package and environment manager. In addition, the pre-processing step also uses the PDAL library (PDAL Contributors, 2020), semantic segmentation uses PyTorch libraries (version 1; Fey and Lenssen 2019) and instance segmentation uses Networkx (Hagberg et al., 2008).

## 3. Results

### 3.1. Plot wide segmentation

A total of 10,557 trees were segmented from 10 plots, of which 3,908 trees were inside the plot boundary and have a *DBH*>0.1 m (Table 3). The tallest tree segmented is a 78.9 m dipterocarp from MLA and the largest tree by volume is a 211 m^3^ coastal redwood from CALI (CALI-A, Figure 3). Trees from WYT have far longer total branch length where the longest is 8.5 km, this compares to the longest tropical branch length of <2 km. This could be as a result of a trees ontogeny e.g. repeated pollarding followed by abandonment, or a systematic method bias where aggressive semantic classification misclassifies smaller branches as leaves for evergreen trees.

**Table 3:** Summary of *TLS2trees* segmented trees, only trees that are within the plot boundary and *DBH* >0.1 m are considered. 99^*th*^ percentile is used to remove outliers.

| Plot | N |  | Height (m) |  |  | Projected crown area (m<sup>2</sup>) |  | Total Volume (m<sup>3</sup>) |  | Trunk volume (m<sup>3</sup>) |  | Branch length (m) |  | DBH (m) |  |
| --- | --- | --- | --- | --- | --- | --- | --- | --- | --- | --- | --- | --- | --- | --- | --- |
|  | \|S\| | S ha<sup>-1</sup> | median | 99<sup>th</sup> | max | median | 99<sup>th</sup> | median | 99<sup>th</sup> | median | 99<sup>th</sup> | median | 99<sup>th</sup> | median | 99<sup>th</sup> |
| RUSH | 162 | 247.0 | 12.5 | 24.5 | 26.1 | 15.8 | 156.7 | 0.3 | 4.2 | 0.2 | 2.0 | 54.0 | 594.8 | 0.3 | 1.3 |
| WYT | 393 | 393.0 | 18.4 | 27.0 | 30.5 | 36.1 | 291.4 | 1.3 | 20.9 | 0.5 | 5.2 | 337.7 | 3570.6 | 0.3 | 1.0 |
| NOU | 547 | 521.0 | 17.9 | 46.0 | 58.9 | 36.6 | 338.7 | 0.4 | 15.9 | 0.3 | 11.7 | 35.8 | 605.3 | 0.2 | 1.2 |
| KOG | 292 | 268.9 | 6.4 | 25.7 | 26.7 | 24.2 | 228.6 | 0.4 | 10.3 | 0.2 | 5.4 | 41.6 | 1166.8 | 0.3 | 1.6 |
| CALI | 804 | 326.4 | 23.9 | 68.4 | 72.9 | 51.1 | 283.9 | 2.3 | 85.7 | 1.8 | 75.4 | 111.0 | 1758.0 | 0.5 | 3.0 |
| MLA | 401 | 371.1 | 15.8 | 65.1 | 78.9 | 30.2 | 543.3 | 0.4 | 35.9 | 0.3 | 32.9 | 33.6 | 1066.1 | 0.3 | 1.5 |
| AEP | 462 | 760.7 | 11.5 | 22.5 | 23.4 | 19.8 | 128.1 | 0.2 | 3.3 | 0.1 | 2.4 | 24.3 | 270.7 | 0.2 | 0.9 |

Estimated segmented stem density ranges from 247 stems ha^-1^ at the RUSH plots to 761 stems ha^-1^ at the AEP plot (Table 3). Compared to reported stem density values (Table 1) these are ±10% for RUSH, WYT, NOU and MLA; larger discrepancies are evident at AEP (−25%), KOG (+39%). Fujimori (1977) reported a large interval for stem density in CALI-type (coastal redwood) plots as a function of succession; values reported here fall within that range.

Figure 4 shows per plot the largest segmented tree point clouds (by total woody volume); for each tree, both wood and leaf points are displayed (and coloured accordingly). Appendix B presents all trees segmented for plots where *TreeQSM* computed *DBH* >0.1 m. In general, tree crowns appear complete with only small omission and commission errors evident e.g. small twigs/branches from neighbouring trees. The crowns of large tropical trees (plots MLA and NOU) are well segmented, even capturing the idiosyncrasies of crown morphology e.g. tree MLA-Q has broken and repsprouted (top row Figure 4). Along the length of some tree stems there are erroneous leaf points that are either attributable to lianas or crowns of smaller mid and understorey trees (e.g. trees NOU-A and MLA-B in Figure 4). As described in Section 2.2.3, leaf points *P_L_* are connected to a stem via branch tips, this would suggest that graph connections have been made to branch points surrounding the stem. It should be noted that *P_L_* are not used to model volume.

**Figure 4:**
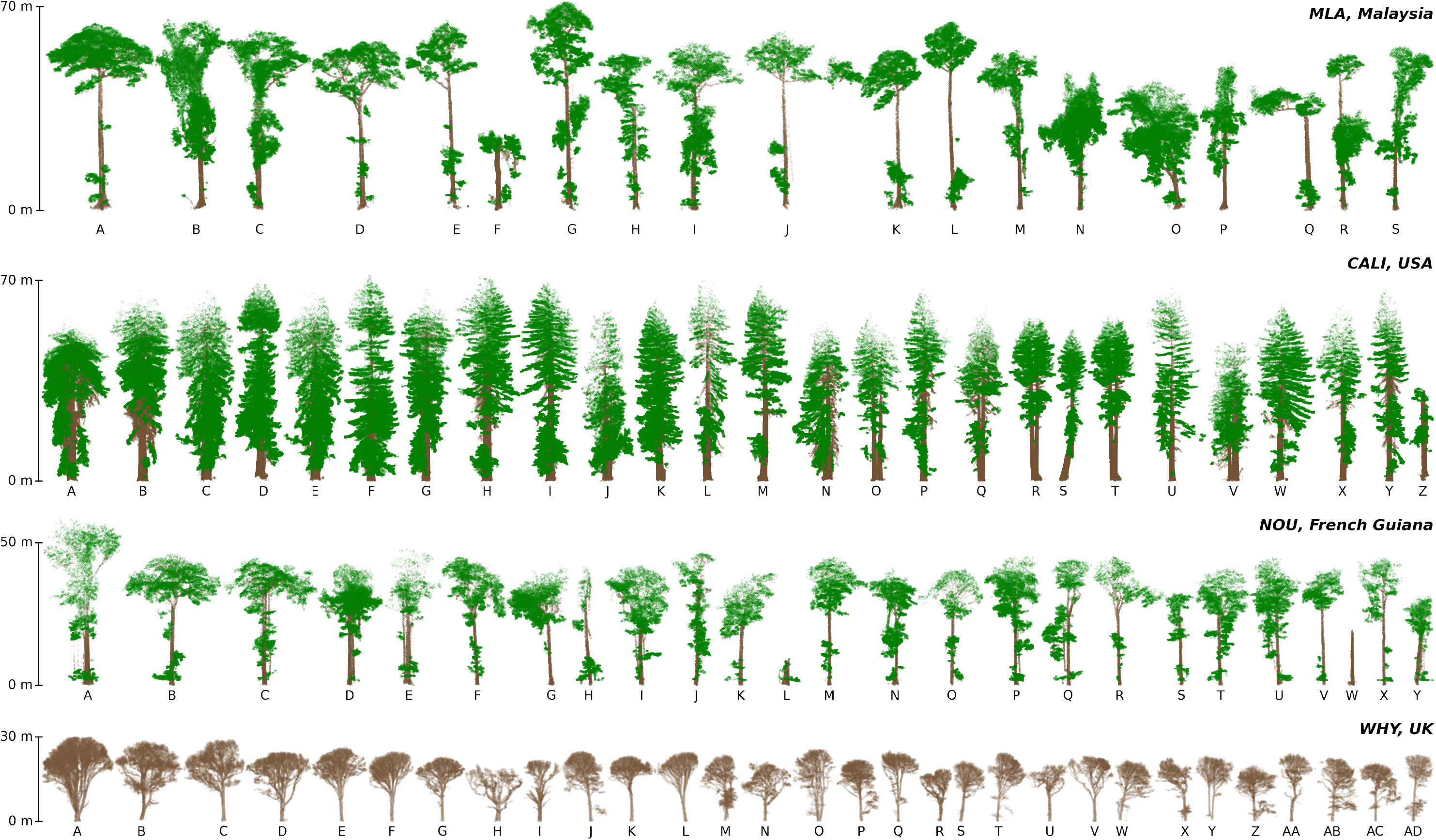
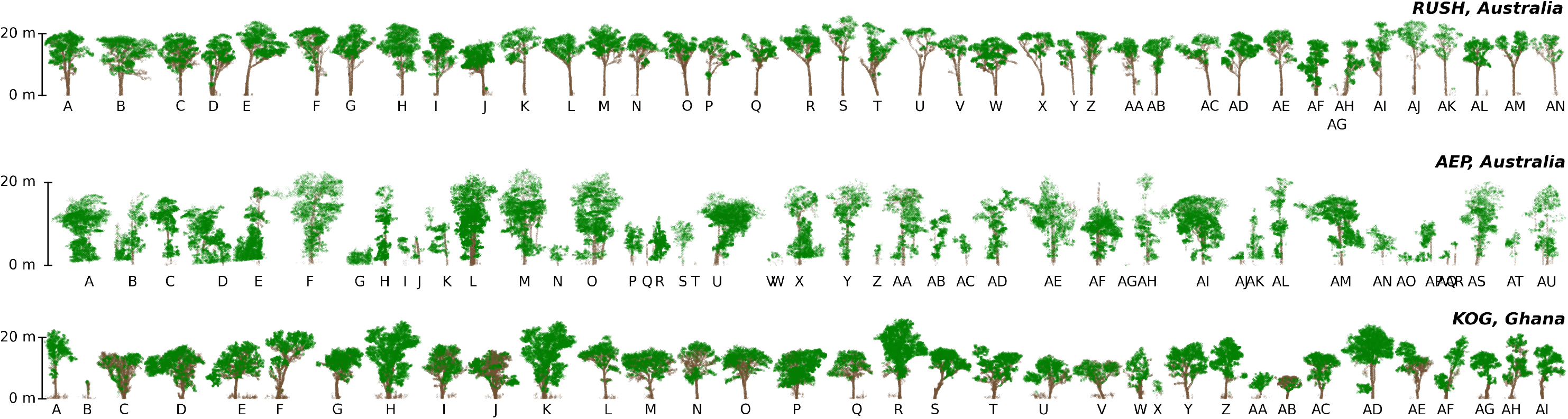
The largest trees (by volume) segmented using *TLS2trees* for different forest types. Points are coloured leaf or wood components as classified during the semantic segmentation step.

*TreeQSM* appears to have overestimated the total volume of a few trees (e.g. trees KOG-B, RUSH-AG, AEP-H, AEP-J and NOU-L in Figure 4). This is a result of commission errors that have inflated the size of cylinders used to estimate volume, particularly at the base. Further, a number of trees in the KOG plot have numerous smaller stems at the base of each tree (see Section 4).

### 3.2. Comparison with manually segmented trees

A total of 1,053 segmented trees were successfully paired to a manually segmented tree, this represents a match rate of 82% (Table 4). For the majority of plots a corresponding tree pair was found for >90% of trees and for all plots a pair was found for all segmented trees that constitute the largest 50% of total segmented volume. The lowest rates are observed at WYT, see Section 4 for discussion.

Scatter plots that compare key QSM and point cloud metrics are presented in Figure 5. Considering total woody volume (a precursor for estimating AGB) *TLS2trees* derived estimates has a per plot bias of ±0.4 m^3^ and RMSE of 0.5-3 m^3^. The exception being CALI where bias is −1.4 m^3^ and a woody volume RMSE of 16.7 m^3^ (%RMSE=70%). Segmentation of smaller trees by *TLS2trees* was less accurate and this results in a per tree total woody volume %RMSE of >1000%. However, when considering the largest trees that constitute 50% of total woody volume, %RMSE of per-tree woody volume reduces to ^~^60%.

**Figure 5:**
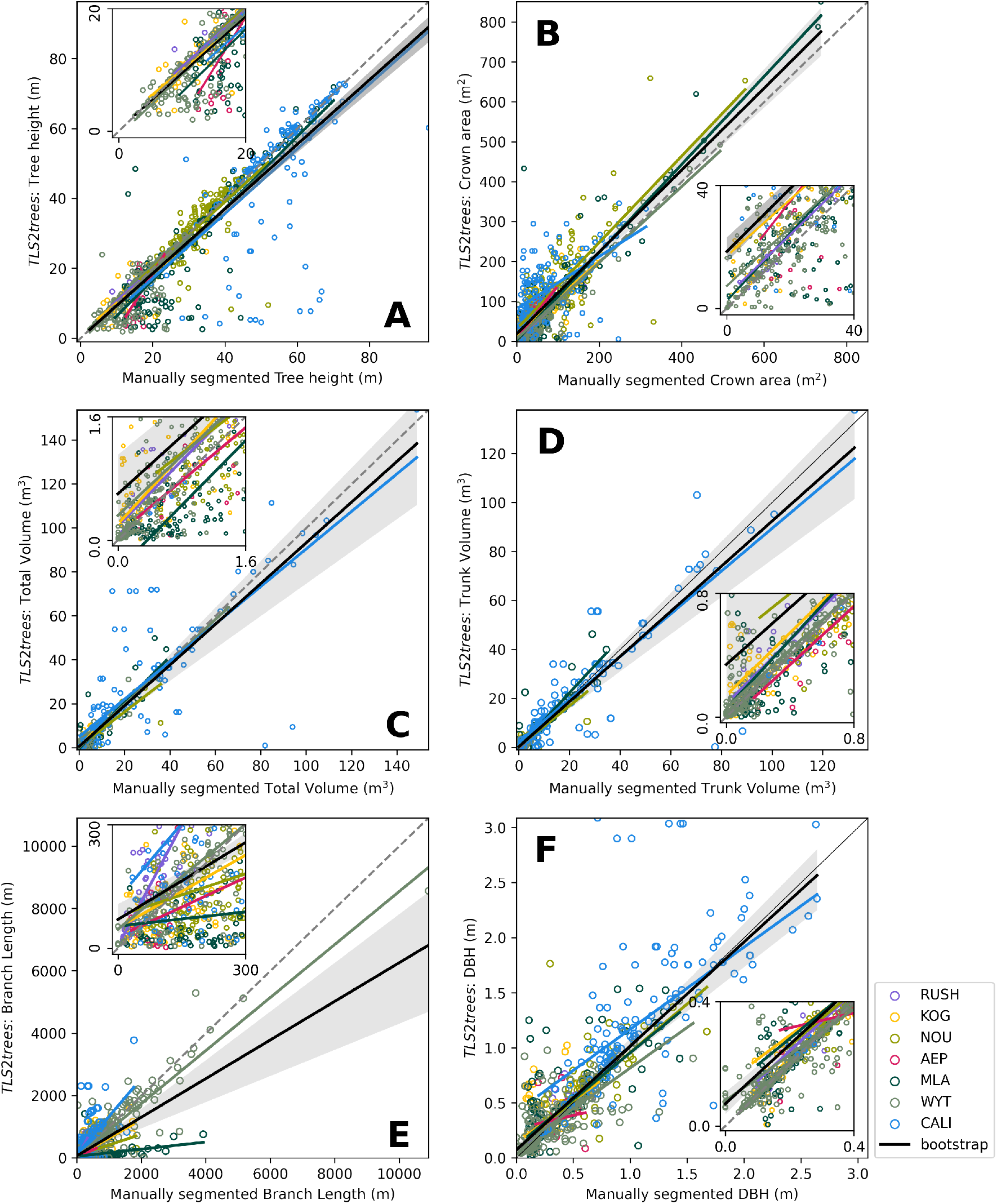
A comparison of manually and *TLS2trees* segmented tree pairs for metrics (A) tree height, (B) crown area, (C) total volume, (D) trunk volume (E) total branch length and (F) *DBH*. Smaller panels are zoomed into the lower 50th percentile of tree pairs. The “boostrap” line represents a mean regression line for all plots, the grey shaded area is a 99% confidence interval for the regression.

A comparison of per plot summed total volume for manually and *TLS2trees* segmented trees shows *TLS2trees* derived total volume were ±10% that of manually segmented trees for all plots (Figure 6). The exception was the RUSH and KOG plots where an 18% and 30% overestimate in total woody volume is evident.

**Figure 6:**
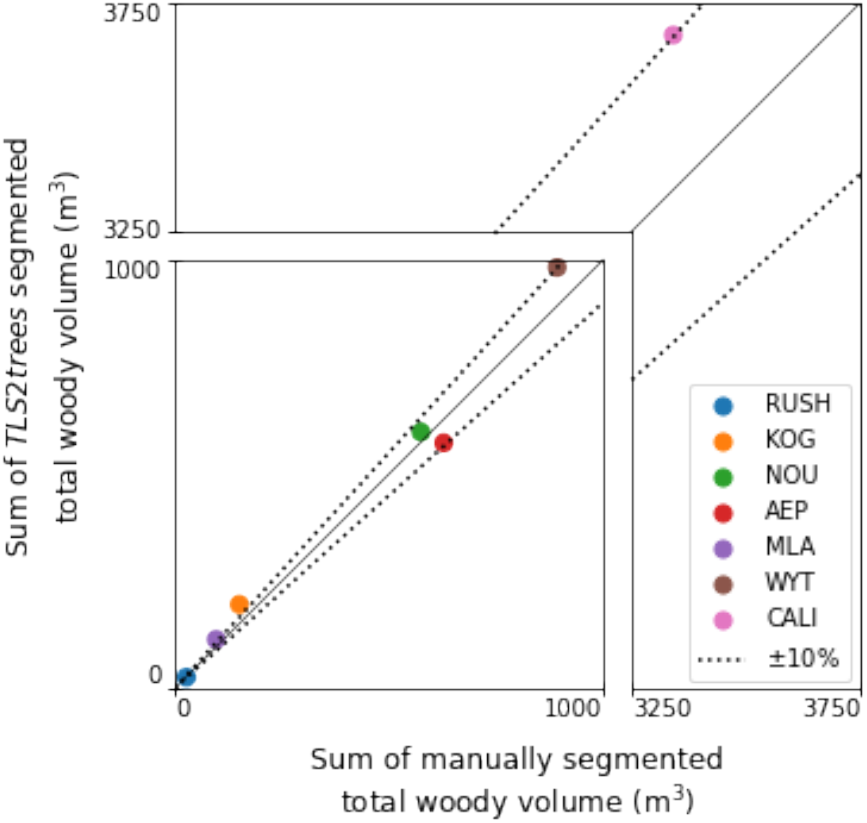
A comparison of per plot summed woody volume for manually and *TLS2trees* segmented trees.

If using an existing allometric equation to estimate tree volume or AGB then other structure metrics are important e.g. *DBH* and tree height. Tree height is estimated with an %RMSE of 27% where the largest errors were at the tall tree sites (MLA and CALI) as well as AEP. For AEP, tree height was underestimated owing to semantic segmentation errors near the base of trees where wood points were classified as leaf, this also impacted estimates of *DBH*. Crown area was consistently overestimated by *TLS2trees* with a bias of 20 m^2^. *DBH* was also overestimated by *TLS2trees* with a bias of 0.04 m and an %RMSE of ^~^ 100%, this reduced to <40% when considering the cohort of largest trees. Inflation of crown area and *DBH* by *TLS2trees* are attributed to outlier points that increase projected area; it is suggested that a method to filter outlier points could improve results.

Segmented tree pair similarity was tested using the Jaccard Index (see Section 2.5). Mean Jaccard Index results for all trees was 75%; this indicates that a pair of trees shared 75% of the voxel space (weighted by point density; Table 4). Differences were predominantly caused by the foliage around stems (e.g. top row Figure 4). Considering all points, a Jaccard Index of >90% is evident at 3 the least densely stocked plots the sites. Jaccard index values increase for all plots when considering only the largest trees that comprise 50% of total woody volume and are generally greater than 75% (except CALI plots; Table 4). Trees at AEP performed poorly when considering Jaccard Index values, with a median score of 13% increasing to 75% when considering the 7 largest trees; it is suggested this is again due to a poor semantic segmentation.

Considering wood and leaf point voxel occupancy (Table 4), the Jaccard Index values were lower than for all-points values. This indicates that there is a mismatch between wood/leaf point segmentation methods. For example at MLA 67% of points were classified as leaf using *TLSeparation* whereas 91% of points were classified as leaf using the deep-learning segmentation (Table 4). This also impacts total branch length (Figure 5E) where *TLS2trees* significantly underestimates length compared to manually segmented trees.

**Table 4:**
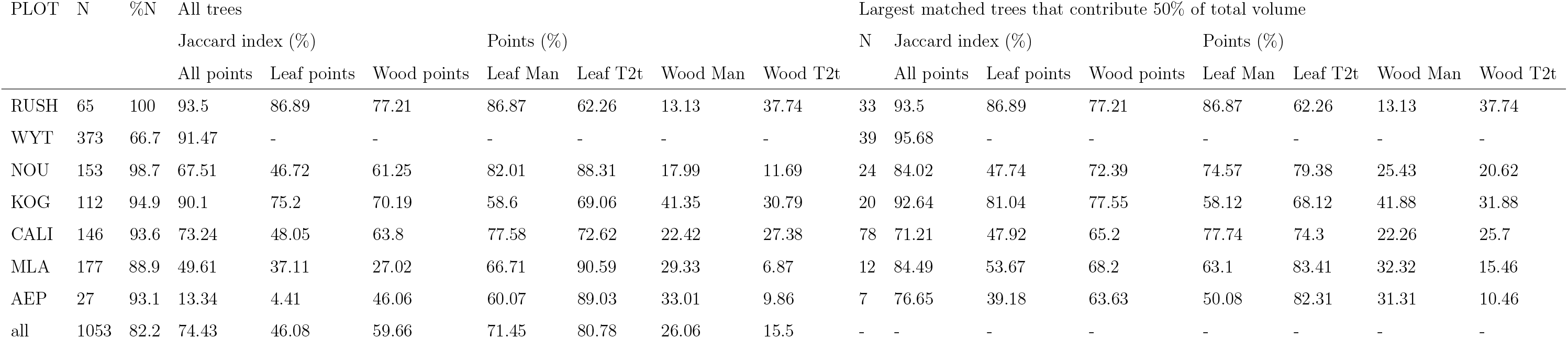
Matched tree pair point clouds comparison between manually and *TLS2trees* (T2t) segmented trees.

## 4. Discussion

TLS methods are capable of generating accurate estimates of tree and forest AGB. Operationalising TLS workflows could therefore have implications for activities including National Forest Inventories (NFI) (Liang et al., 2018), Measurement, Reporting, and Verification (MRV) protocols, benchmarks and reference data sets for AGB focused satellite missions (e.g. BIOMASS and GEDI) (Chave et al., 2019) and new or updating allometry (Stovall et al., 2017, Disney et al., 2020). However, achieving operationalisation of TLS methods has proved challenging for a number of reasons.

In particular, and the focus of this work, is the labour-intensive and time-consuming efforts required to accurately segment individual trees from plot-level point clouds which has caused a significant processing bottleneck. To address this issue, here we have presented *TLS2trees*, a FOSS and horizontally scalable Python-based pipeline for segmenting individual trees from plot-scale TLS point clouds.

Manually segmenting trees from plot-level point clouds is currently regarded as the most accurate and, it is suggested, that this is unlikely to change. However, such approaches are not reproducible and lack the traceability or transparency that would be required for carbon accounting and reporting schemes. Manual segmentation is also not scalable, where, along with issues of subjectivity, operator fatigue may degrade results. An automated approach, such as *TLS2trees*, aims to address these issues of reproducibility, subjectivity and scalability. However, routinely segmenting many hundreds of trees per hectare presents additional challenges, for example, quality assurance (QA). As seen here, different forest types present different challenges in terms of generating an accurate result and therefore an application focused QA strategy is suggested. For example, if the metric of interest is AGB, a strategy of sampling the largest trees, that disproportionately contribute to plot level AGB, as well as a subset of smaller trees across the AGB range should identify systematic issues.

*TLS2trees* performed well across a range of forest types, from open savanna (KOG) to tall tropical rainforest (MLA), the workflow also performed well within plots segmenting trees across a range of sizes (Table 3). This is despite the fact the semantic segmentation base model was only trained on a small area of Australian and New Zealand forest (Krisanski et al., 2021) and *TLS2trees* model parameters were not adjusted for forest type (Appendix A).

We would also like to stress that, like all software packages, *TLS2trees* is a work in progress and there are a number of aspects of *TLS2trees* that could be improved. We hope this can be achieved through a user community and have made the source code open-source to facilitate this. The success of each step presented in Figure 3 depends on the success of the previous step. For example, as seen at the AEP plot, mis-classification of wood points as leaf points around the base of trees caused errors in the instance segmentation. Similarly, QSM methods such as *TreeQSM* require a “clean” point cloud e.g. with minimal noise (Raumonen et al., 2013) and this was not always achieved e.g. small neighbouring stems led to volume inflation. It is suggested that improved results can be achieved by testing model parameterisation on individual tiles before scaling up. Another approach could be to test different parameterisation in a simulated forest, for example a comparison of semantic segmentation methods (Morel et al., 2020).

One particular aspect where *TLS2trees* performed poorly was with the segmentation of smaller trees. Example errors included commission errors where multiple stems were grouped into a single tree or omission errors where smaller stems were missed. Again, a different parameterisation of *TLS2trees* may improve performance. An example is presented in Figure 7 where a change in the cumulative gap parameter reduces the number of erroneous stems at the base of the tree; however, if the parameters is reduced too far then portions of the crowm are removed. Instrument limitations should also to considered when resolving smaller features, where laser beam and scanning characteristics can limit capability (Demol et al., 2022b). An example of the impact of the smallest trees is presented for WYT where total volume %RMSE was ^~^2000% when compared to manually segmented trees. However, if considering only the largest trees that contributed 90% and 50% to total volume, %RMSE reduced to 50% and 25% respectively. Further, matching pairs of trees was a challenge at WYT. Coppiced stools, where trees forked below 1.3 m, were often regarded as a single tree by *TLS2trees* whereas they were manually extracted as individuals by Calders et al. (2018). A suggested solution to improving instance segmentation for small, forked trees is to perform another instance segmentation based upon topological information derived from the QSM.

**Figure 7:**
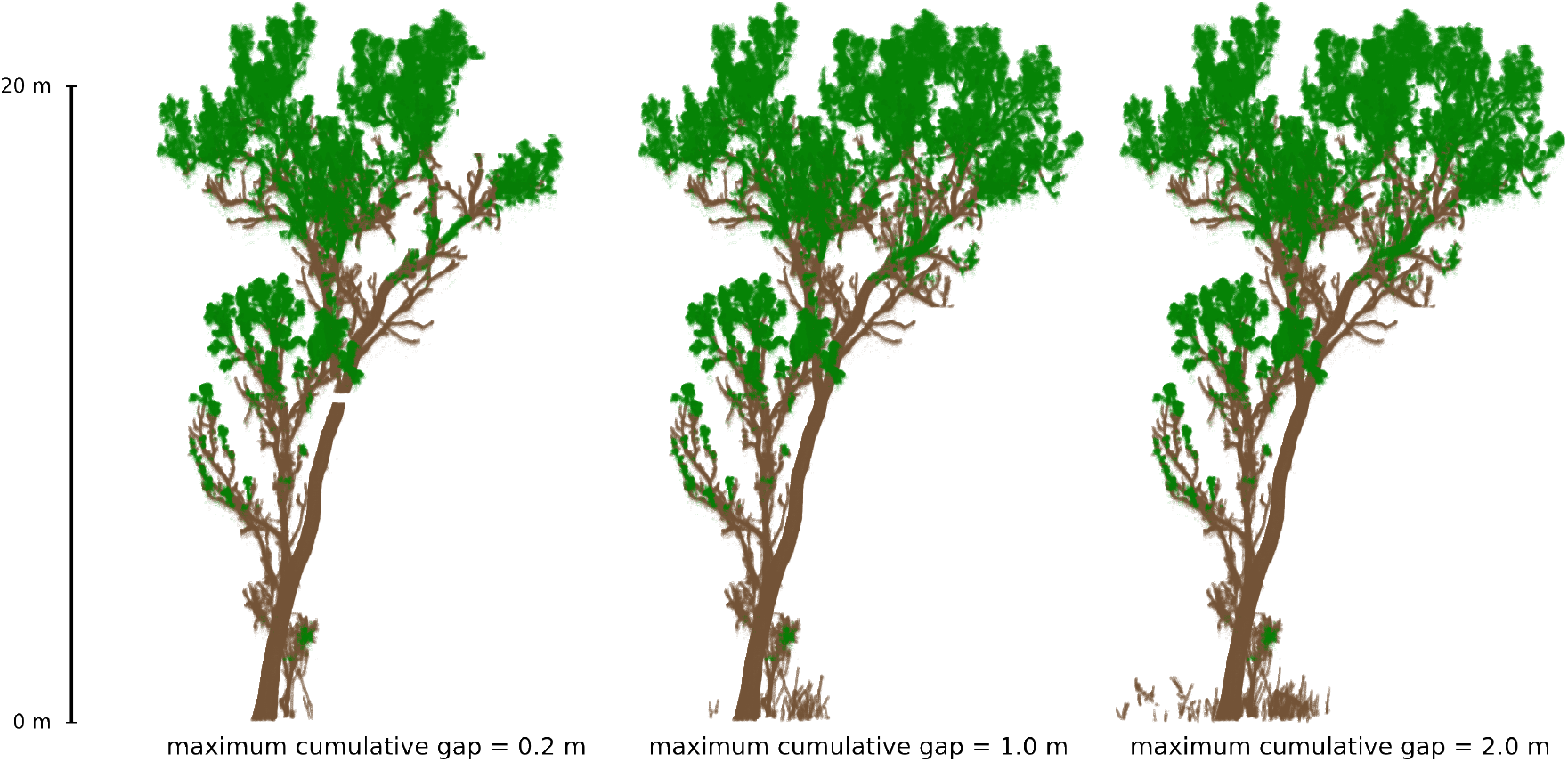
An example of different outcomes when altering model parameters. The parameter `--maximum-cumulative-gap` alters the maximum cumulative gap between node and a base node. In this example, a low value segments the base well but removes smaller branches at the top of the canopy whereas a higher value results in small stems around the base being included. The value used for this manuscript was `--maximum-cumulative-gap`=2 m

Considering the scalability of *TLS2trees* there may be a number of options for improving code and routine efficiency. For example, a 10 m × 10 m processing unit (see Section 2.2.1) may not be optimal for all scenarios and processing architectures, or a larger vertical slice (and therefore reduced number of clusters) in the semantic segmentation step (see Section 2.2.3) may also improve compute times. Here, *TLS2trees* was run on a state-of-the-art computing facility that may not be available to all groups. An alternative option would be to run on a commercially available cloud computing service (e.g. Amazon Web Service or Microsoft Azure), to facilitate this we have provided a containerised version of *TLS2trees*.

Lastly, although the name *TLS2trees* implies the workflow is limited to the processing of just TLS data, we suggest that the framework presented here could be applied to other laser scanning modes. The horizontal scalability functionality, built in to *TLS2trees*, is well suited to the large area acquisitions possible with airborne platforms.

## 5. Conclusion

There is an estimated 260 ha of plot-level TLS data collected from forests across the globe (*pers. comm*. Dr. Atticus Stovall, NASA, 22nd November 2022); processing this data archive could yield upwards of 260,000 individuals trees. The biophysical and ecological insight that could be drawn from this data, including and beyond the estimation of AGB, could be significant. Further, there are a number of other potential uses for this data, including accurate 3D representations of forest plots in radiative transfer (Calders et al., 2018) or other large area modelling approaches. However, much of this data remains unprocessed (to individual tree level) owing to instance segmentation bottlenecks, a problem we suggest is one of the remaining hurdles to operationalising TLS protocols. Here we have presented *TLS2trees*, a workflow and processing pipeline to segment point clouds of individual trees from plot-level TLS point clouds, that we hope will begin to redress this issue.

## Acknowledgments

PW and MD were funded by NERC National Centre for Earth Observation (NCEO). PW, MD, AS, YM and LB were funded under NERC grant NE/P011780/1, MD and AB were funded under NERC grant NE/N00373X/1, YM was funded under NERC grant NE/P012337/1 as well as ERC grant GEM-TRAITS (321121). KC was funded by the European Union (ERC-2021-STG Grant agreement No. 101039795). Views and opinions expressed are however those of the author(s) only and do not necessarily reflect those of the European Union or the European Research Council Executive Agency. Neither the European Union nor the granting authority can be held responsible for them. NO was funded by the MetEOC-4 project which included funding from the EMPIR programme co-financed by the Participating States and from the European Union’s Horizon 2020 research and innovation programme. BB was funded by ESA IDEAS-QA4EO (contract no ESA AO/1-11035/21/I-DT). SK was funded by the Australian Research Council, Training Centre for Forest Value (IC150100004). DC, DM and computing infrastructure were fund by NERC Earth Observation Data Acquisition and Analysis Service (NEODAAS). Capital equipment was funded by NCEO and UCL Geography. Data was collected in Malaysia under permit JKM/MBS.1000-2/2 JLD.7 (87), we are grateful for project partners Sabah Biodiversity Center, Chief Minister’s Department Office of Internal Affairs & Research, Land & Survey Department, Sabah Forestry Department, the Maliau Basin and Danum Valley Management Committees, Forest Research Center (Sabah). We are also grateful for the assistance of Matt Bradford and CSIRO Atherton.

## Author Contributions

PW and MD conceived the project; PW, MD, AB, KC, NO, LD, JA, BB, HB and AS collected data; PW, CCB, BF, AB, KC, BB, HB, DC, DF and NO processed data; PW, SK, DC, DF and WY developed software; PW conducted analysis; all authors contributed to and approved the final manuscript.

## Code and Data Availability

The *TLS2trees* code is available from https://github.com/philwilkes/TLS2trees. Code to run TreeQSM in Octave is available from https://github.com/philwilkes/run-treeqsm-in-octave.

---

Data availability and DOI links

| Plot | Raw data | Segmented trees |
| --- | --- | --- |
| RUSH | LINK | LINK |
| WYT | LINK | LINK |
| NOU | LINK | upon request |
| KOG | upon request | upon request |
| CALI | upon request | upon request |
| MLA | upon request | upon request |
| AEP | upon request | upon request |

