## Appendix 2 for "*TLS2trees*: a scalable tree segmentation pipeline for TLS data"

### Appendix B. Segmented trees

All trees segmented from 10 plots where trees are within the plot boundary and have a *DBH*>0.1 m as calculated by TreeQSM.

*RUSH06*

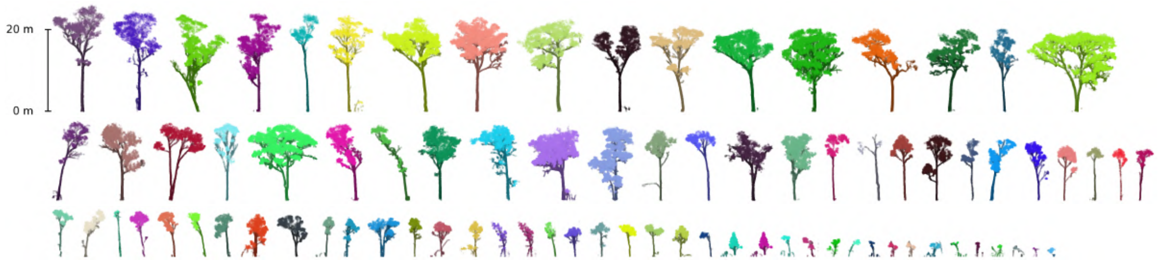


*RUSH07*

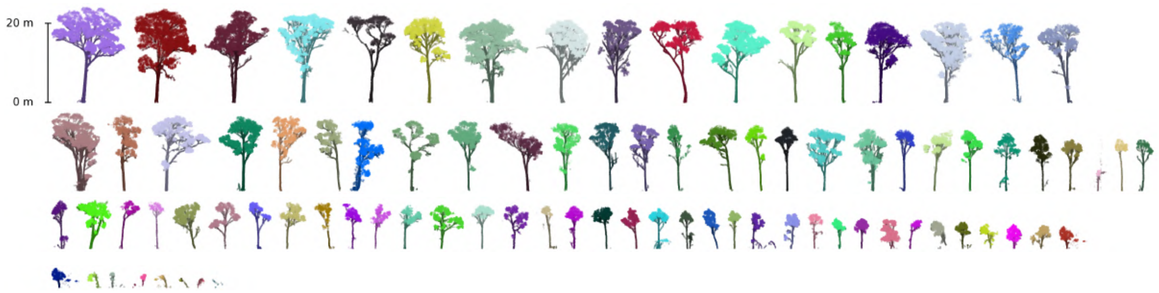


*WYT*

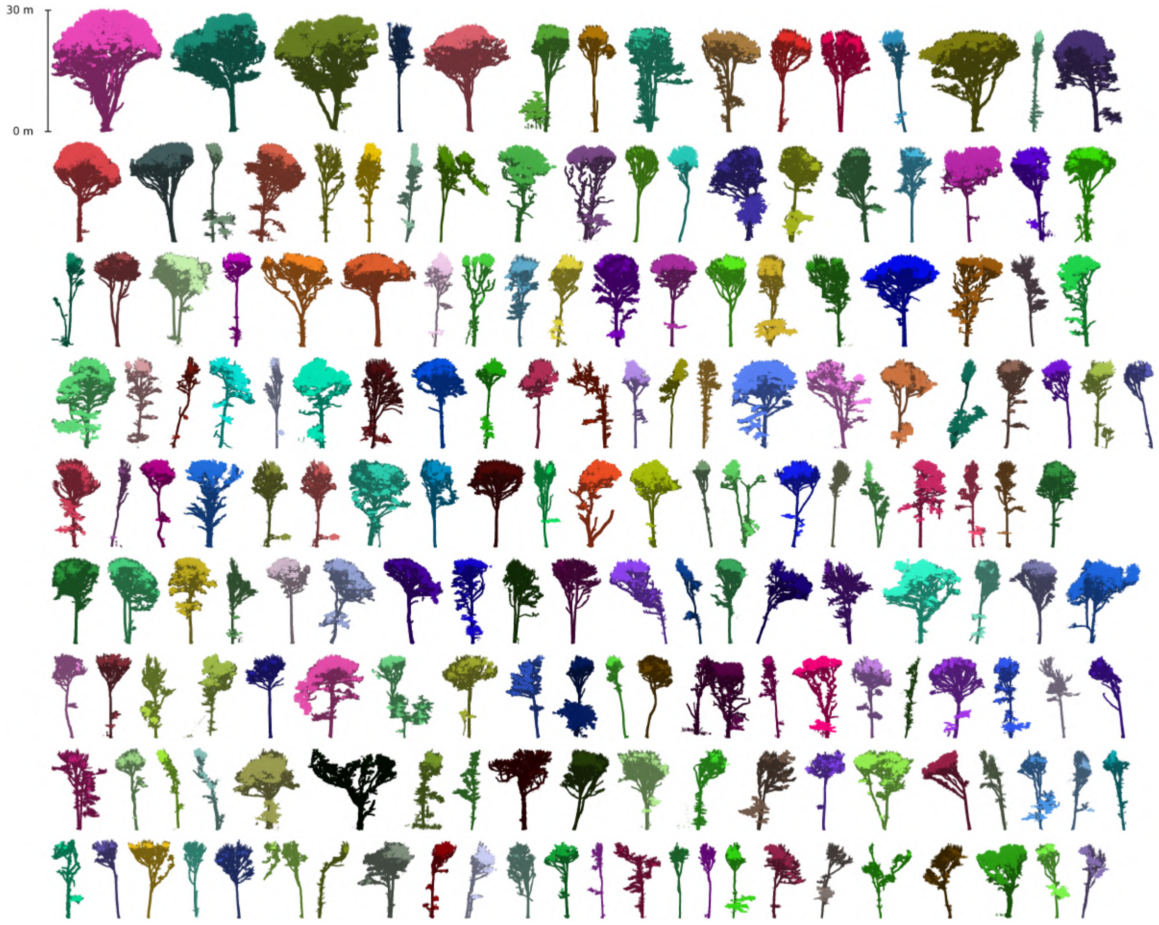

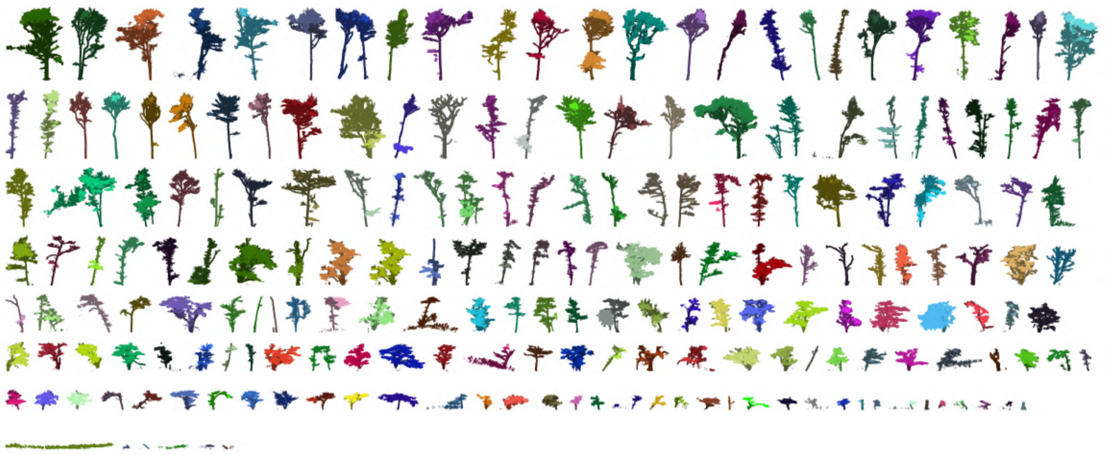

662 *NOU*

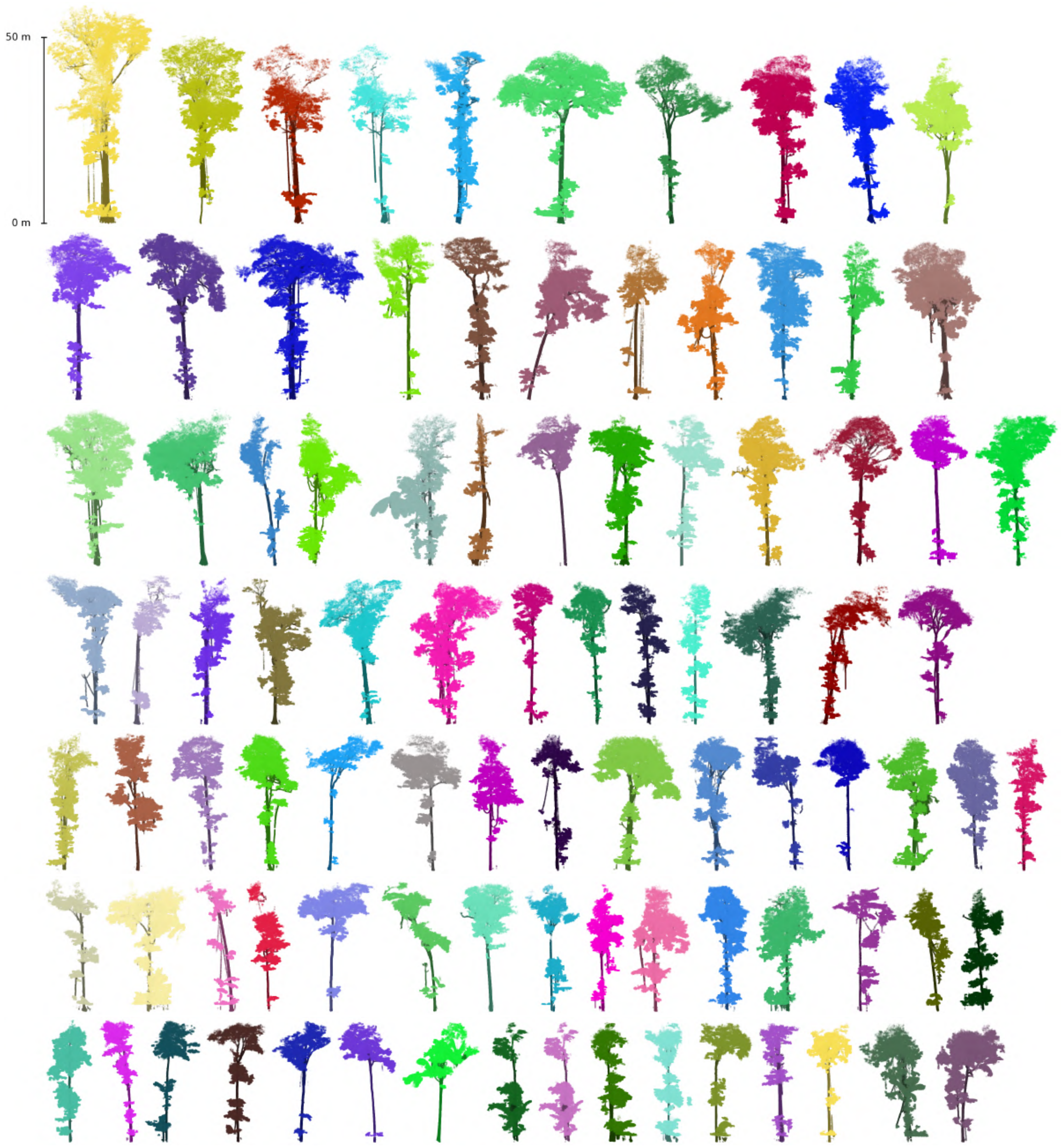

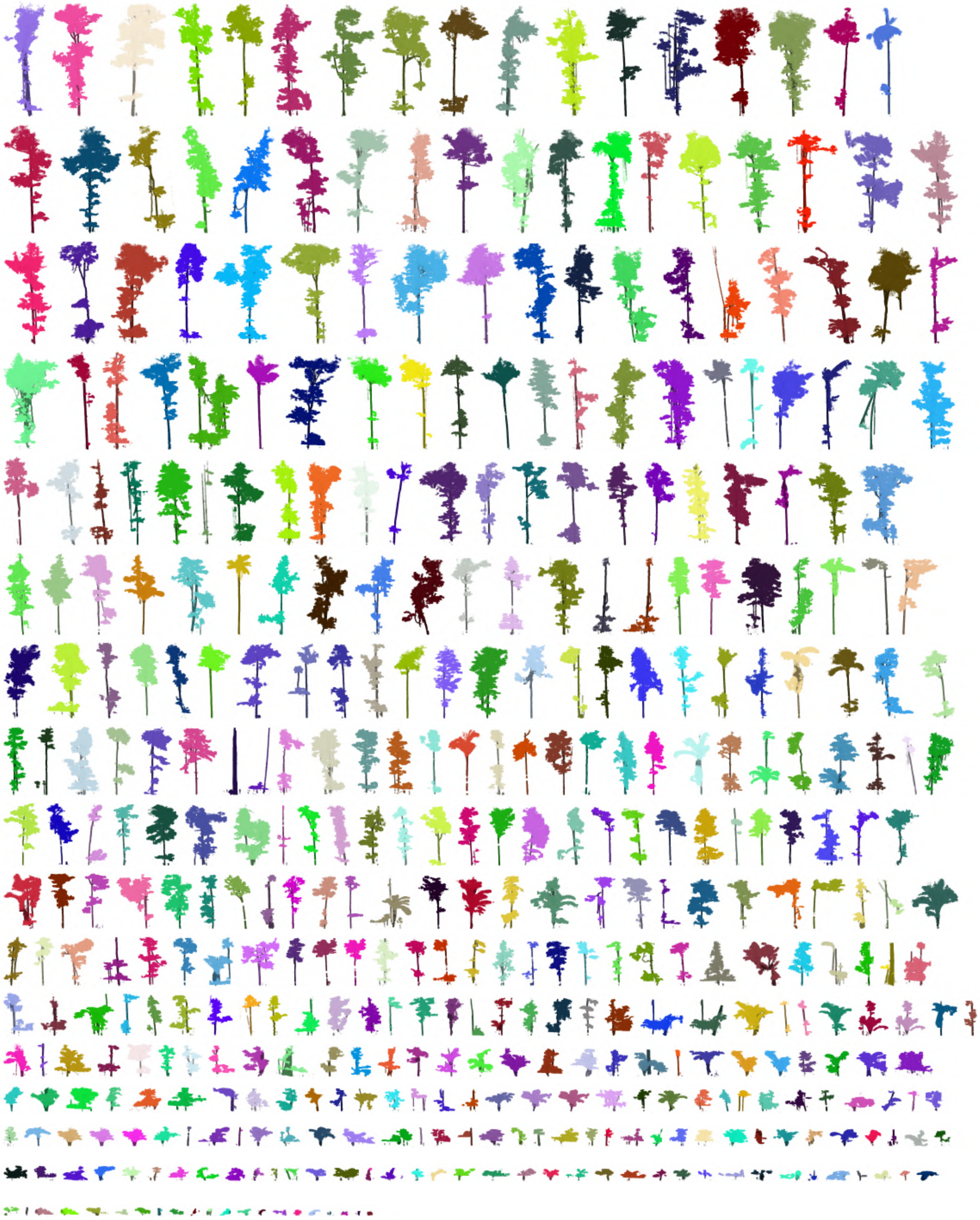

*KOG*

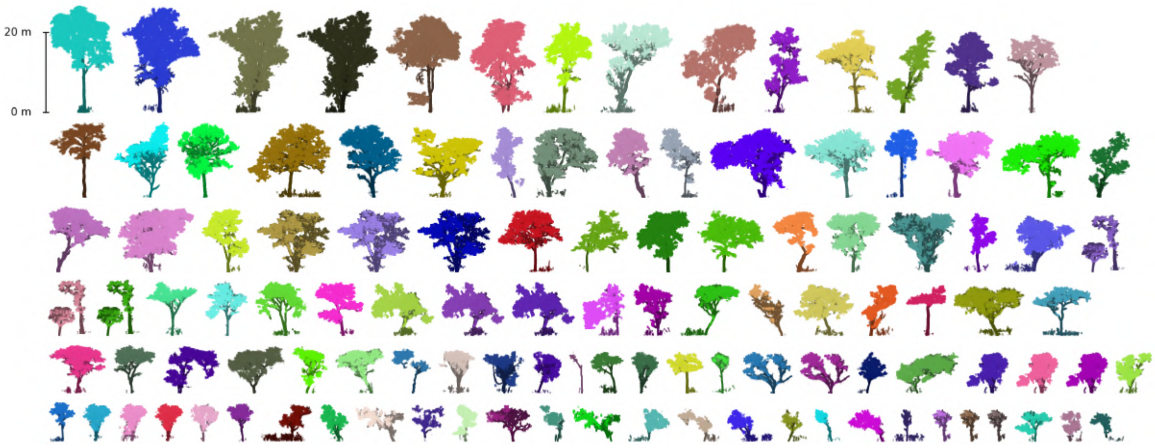

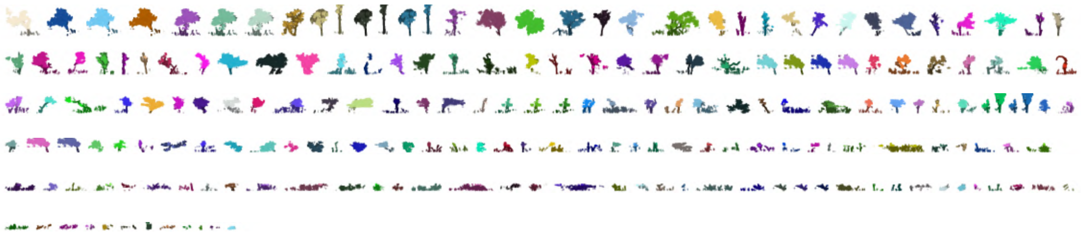

664 *CALI-01*

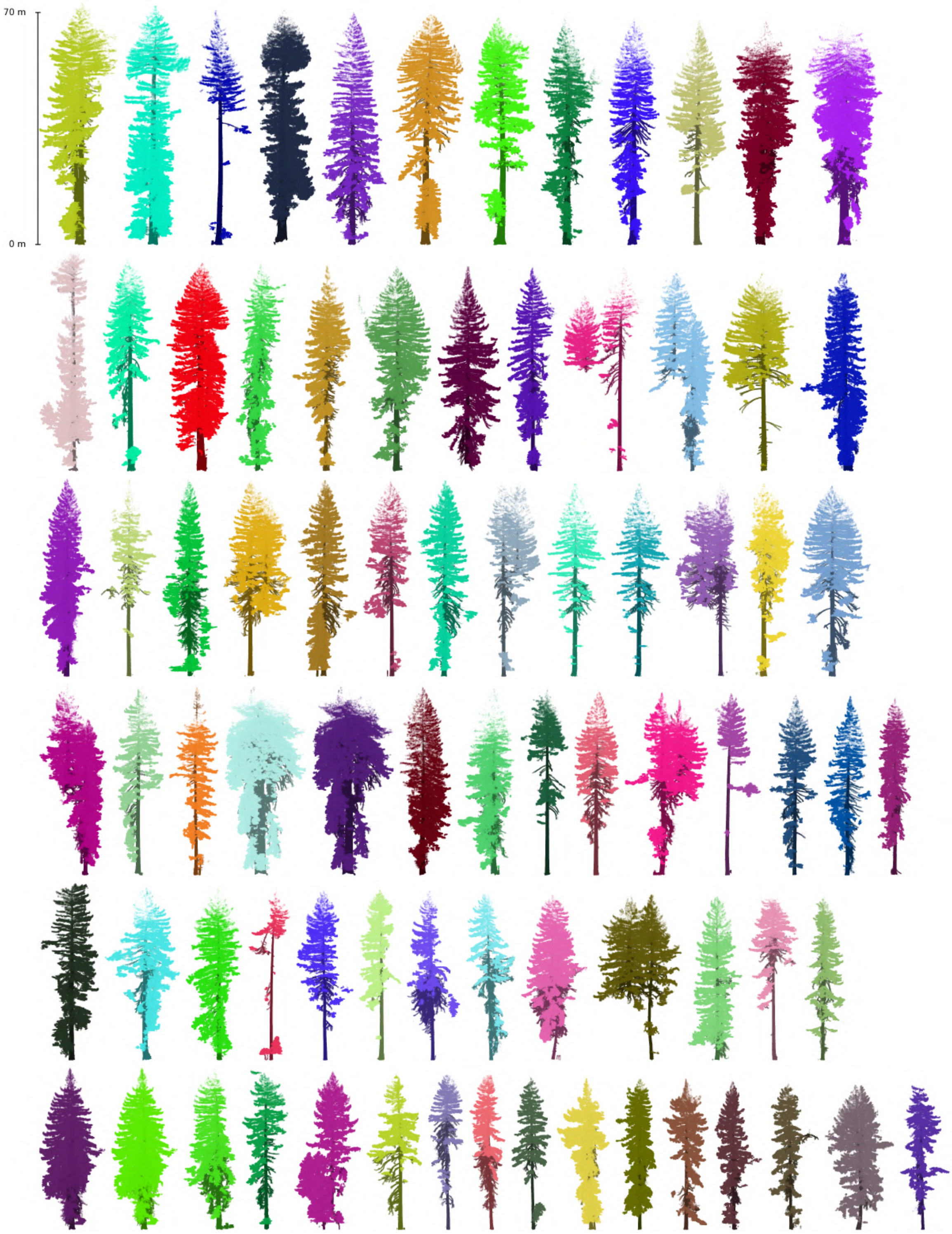

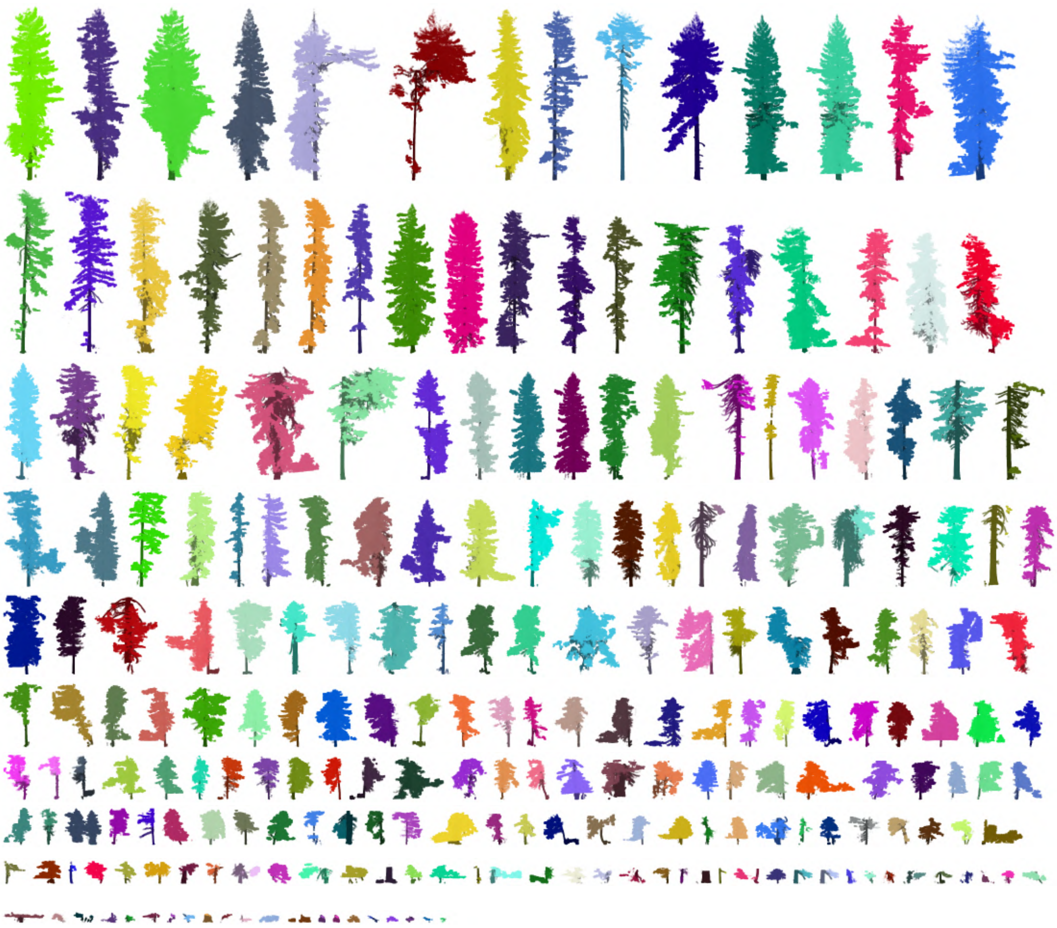

*CALI-02*

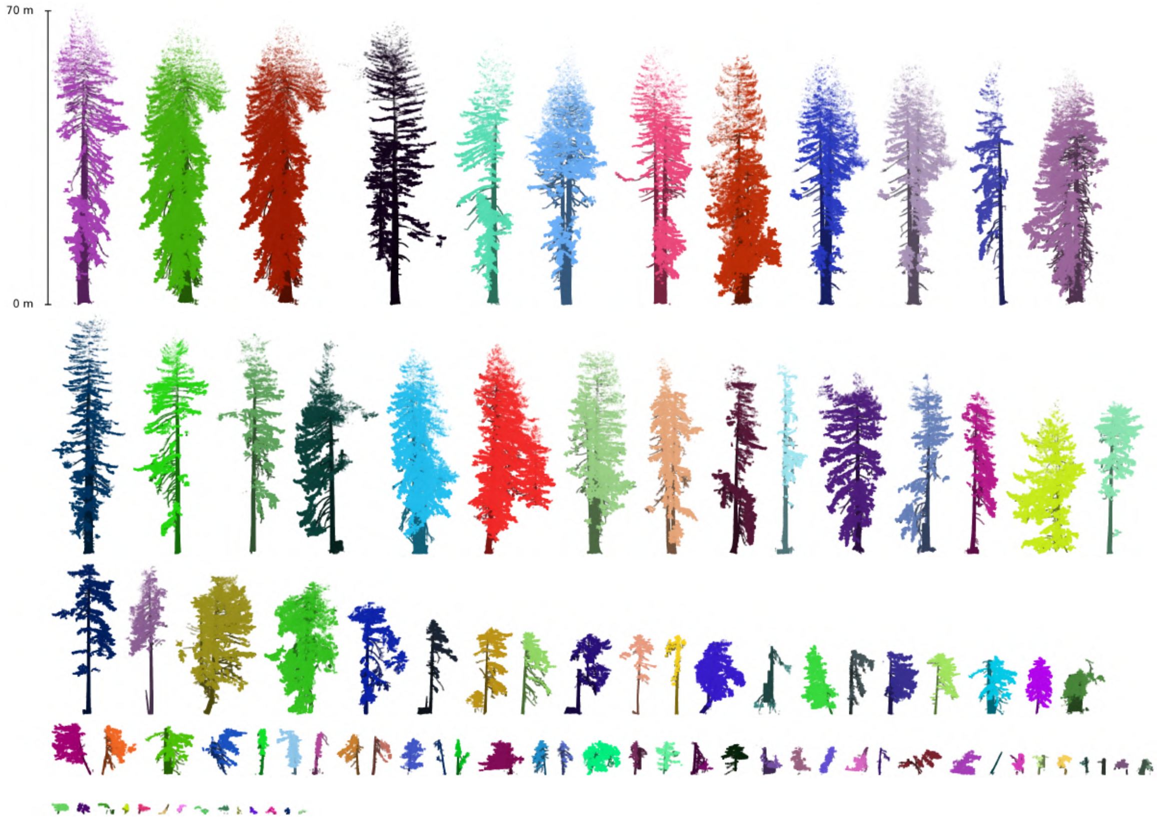

666 *CALI-07*

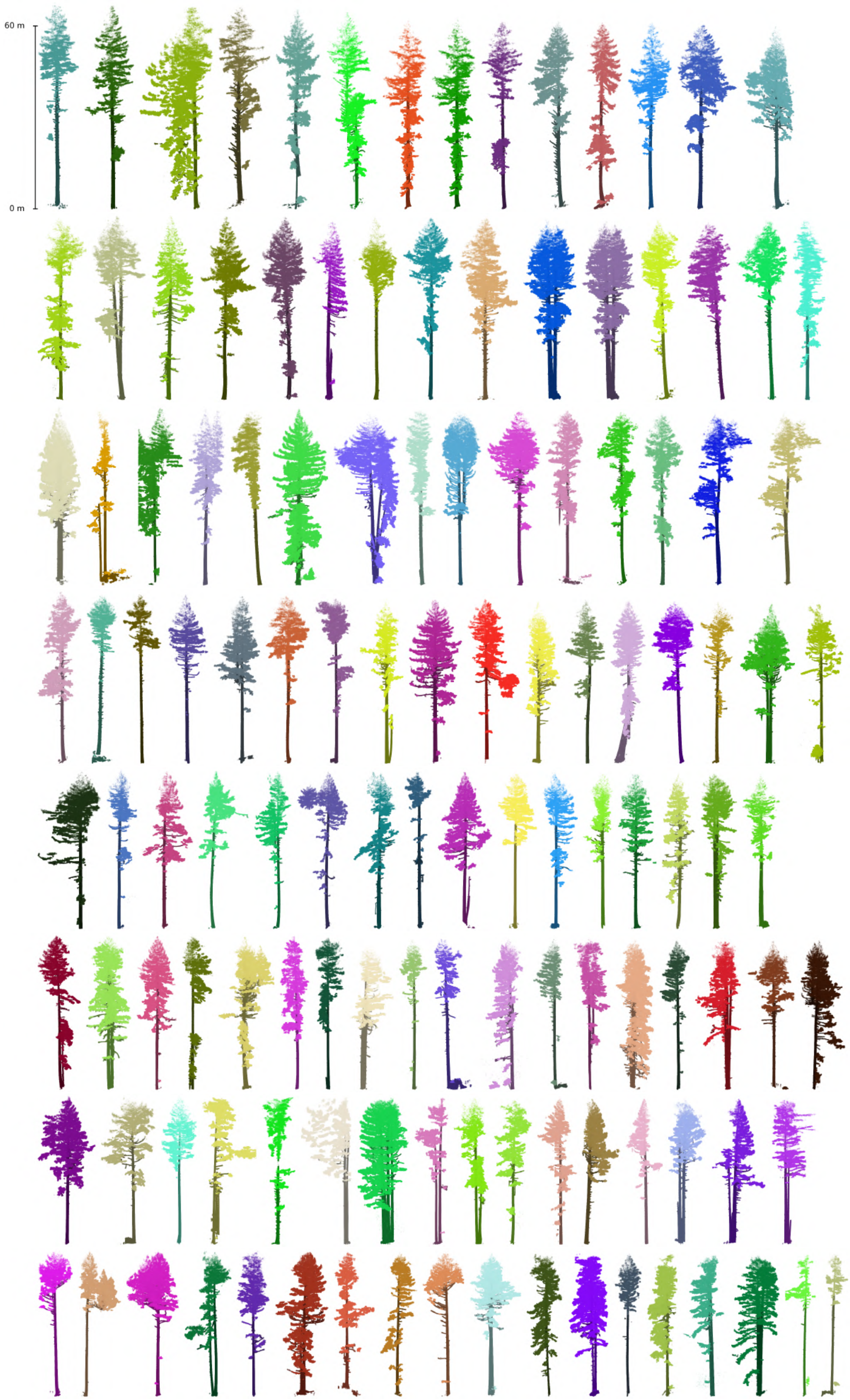

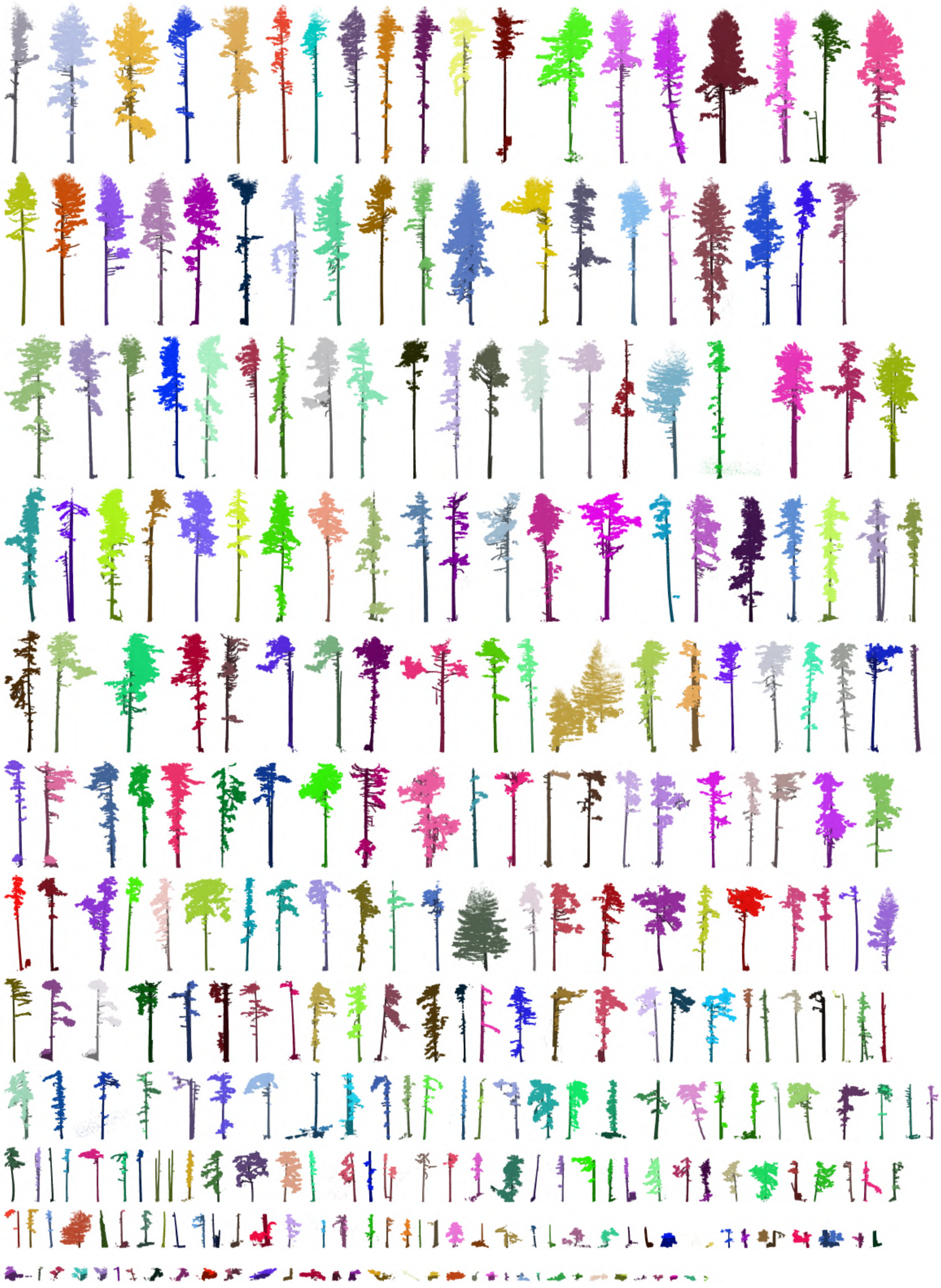

667 *MLA*

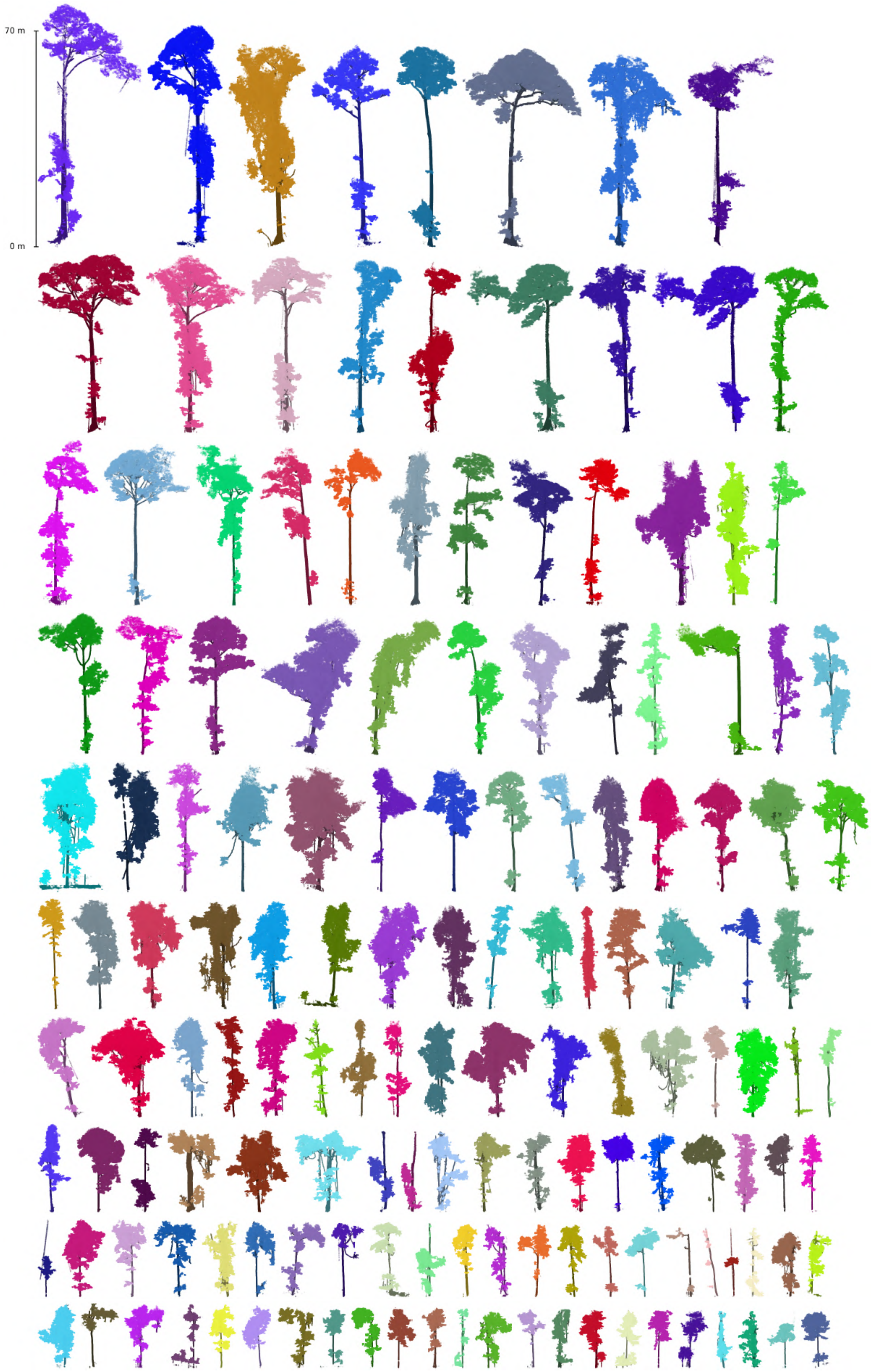

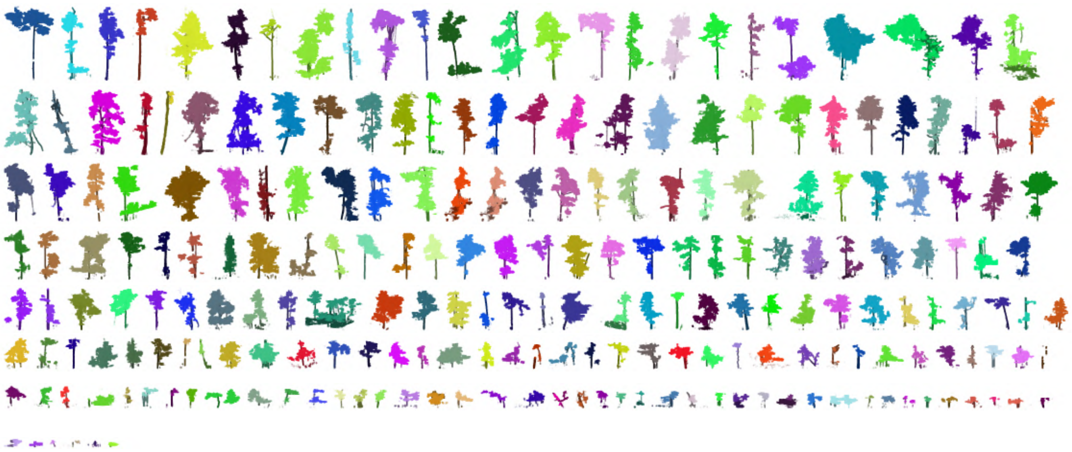

*AEP*

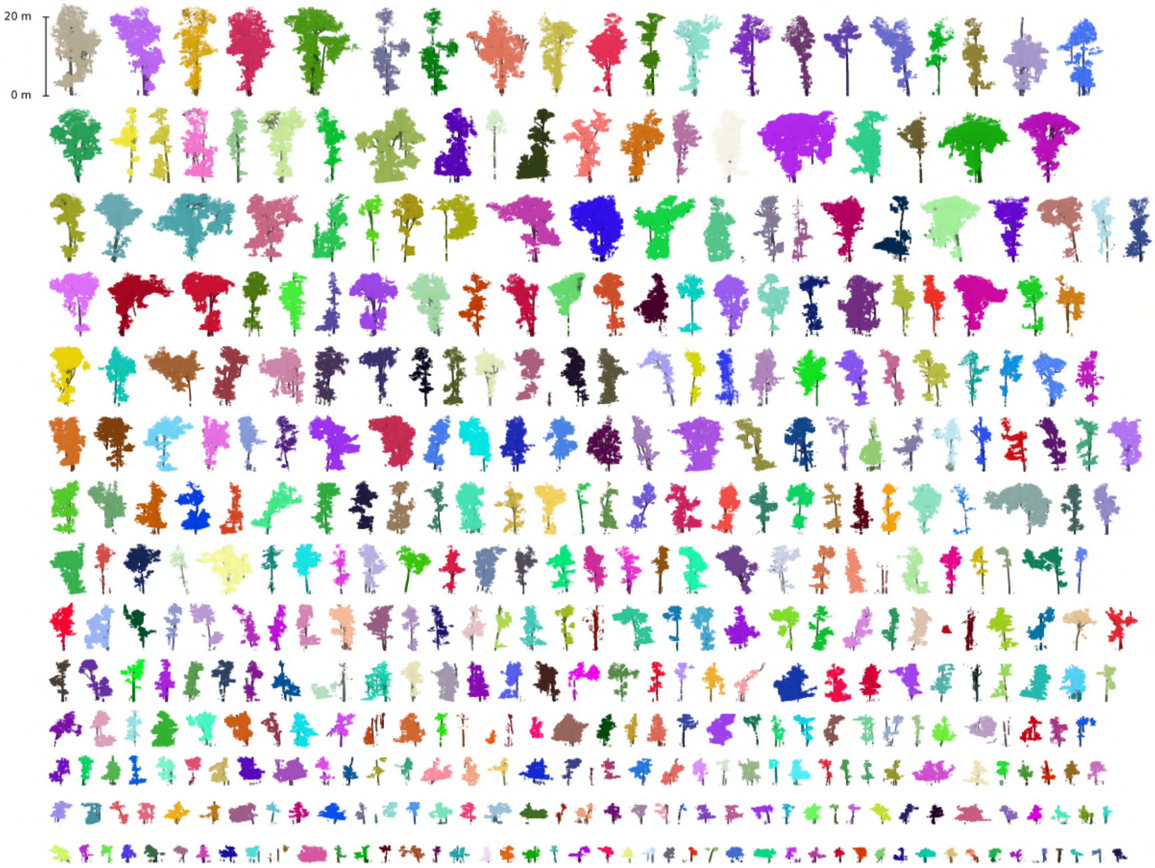
