## Appendix 1 for "*TLS2trees*: a scalable tree segmentation pipeline for TLS data"

### Appendix A. Python tools and exposed parameters

Table A.1: Python commands and parameter values used to segment trees. All code can be found at https://github.com/philwilkes/TLS2trees/. *specific to RIEGL VZ scanners

| Parameter | Purpose | Value used |
|---|---|---|
| *1. Preprocessing* | | |
| **`rxp2ply.py`*** | | |
| `--tile` | Edge length of tile | 10 |
| `--deviation` | Upper and lower bounds of deviation values | 0, 15 |
| `--reflectance` | Upper and lower bounds of reflectance values | -20, 0 |
| **`downsample.py`** | | |
| `--length` | Edge length voxel where 1 point is retained per voxel | 0.02 |
| **`tile_index.py`** | | |
| *2. Semantic segmentation* | | |
| **`run.py`** | | |
| `--buffer` | Size of buffer to use from surrounding tiles | 5 |
| *3. Instance segmentation* | | |
| **`points2trees.py`** | | |
| `--n-tiles` | Number of tiles to use as a buffer i.e. by 3x3 or tiles or 5x5 tiles | 3, 5 or 7 |
| `--slice-thickness` | Vertical slice thickness for constructing graph | 0.2 |
| `--find-stems-height` | Base height for identifying stems | 2 |
| `--find-stems-thickness` | Thickness of slice used for identifying stems | 0.5 |
| `--find-stems-min-radius` | Minimum radius of found stems | 0.025 |
| `--find-stems-min-points` | Minimum number of points for found stems | 200 |
| `--graph-edge-length` | Maximum distance between individual nodes in a graph | 2 |
| `--graph-maximum-cumulative-gap` | Maximum cumulative distance between a base and a node | 3 |
| `--min-points-per-tree` | Minimum number of points for a tree to be segmented | 200 |
| `--add-leaves` | Whether to add leaf points | TRUE |
| `--add-leaves-voxel-length` | Voxel size when add leaves | 0.5 |
| `--add-leaves-edge-length` | Maximum distance used to connect points in leaf graph | 1 |
